# Microchimerism in the Human Brain, Quantitative Assessment and Single Nucleus Profiling Establish Cell Types and Diversity

**DOI:** 10.64898/2026.06.05.730225

**Authors:** Sami B. Kanaan, Ashley McDonough, Coline Gentil, Jeff Ojemann, Haynes Heaton, Reza Behboudi, Dan T. A. Eisenberg, Scott N. Furlan, Dan Geraghty, Joe Rutledge, Francesca Urselli, Jonathan R. Weinstein, J. Lee Nelson

## Abstract

Bi-directional maternal-fetal exchange during pregnancy creates a long-term microchimerism legacy in both individuals, but its presence and cellular fate in the human brain remain poorly defined. We identified maternal microchimerism in the majority of surgically resected epilepsy brains by polymorphism-specific qPCR, across temporal, frontal, parietal, and hippocampal regions. We conducted single nucleus RNA sequencing (snRNA-seq) and analysis by *cellector,* a genetic demultiplexing tool to detect rare allogeneic cells, to test epilepsy specimens and healthy brains from gestation to old age from publicly available snRNA-seq datasets. Microchimerism was detected in both cohorts and revealed diverse neuronal, glial, immune, and vascular identities. L2/3 intratelencephalic neuronal subtypes and microglial/perivascular macrophage were dominant profiles. Microchimerism showed heterogeneous transcriptional profiles and differences from host nuclei. Microchimerism persisted into advanced age but with diminishing frequency. Together, these findings establish microchimerism with cellular diversity as a common feature of the human brain, prenatal through advanced age.

## Introduction

Bidirectional exchange of cells between the mother and fetus during pregnancy creates a life-long legacy of microchimerism (**Mc**) in respective individuals^1,2^. In addition to leukocyte subsets, these semi-allogeneic cells integrate broadly into organs of the recipient and are found with tissue-specific phenotypes including cardiac myocytes, thyrocytes, β-islet cells, hepatocytes, keratinocytes, and endothelial and epithelial cells in various organs^3–8,1,2^. Minimal data is available regarding maternal Mc in the human brain. One study detected maternal Mc in two of three second trimester fetal brain tissues by semi-quantitative PCR^9^. Another study of postmortem brains from six males with multiple sclerosis and six without neurologic disease reported female cells (presumed maternal) in five and six cases, respectively, with immune and neural phenotypes identified in a few^10^. Postmortem brains from females who had Alzheimer’s disease or were cognitively and pathologically normal had Mc with male DNA in the majority, presumed from prior pregnancy with a male fetus^11^. Male Mc in females has been found within major regions of the central nervous system (CNS) including the telencephalon, diencephalon, rhombencephalon, and spinal cord in healthy and diseased subjects^11,12^.

In experimental models, microchimeric cells of maternal and fetal origin have been reported to integrate into the brain and acquire neural phenotypes^13–15^. Interestingly, a recent murine study reported effects of maternal Mc on early fetal neurodevelopment, brain homeostasis, synaptic pruning, and maturation of behavioral abilities^15^. These experimental studies lend credence to strategies exploring chimeric microglia replacement therapies for some types of brain disease^16,17^.

The human brain is seldom biopsied posing challenge to Mc investigation as participation of family members is needed to determine Mc origin and for autopsy specimens family are often not available or approachable. To overcome this challenge, we obtained surgically resected brain tissue from medication-refractory pediatric epilepsy patients and requested mothers and other family members to donate buccal swab samples to genotype for non-shared polymorphisms. We employed a panel of polymorphism-specific highly sensitive quantitative PCR (**qPCR**) assays we developed to interrogate DNA extracted from brain specimens for maternal-specific Mc. The qPCR assays targeted 40 different polymorphisms, most in the polymorphic human leukocyte antigen (**HLA**) region^18,19^. Next, we conducted single-nucleus RNA sequencing (**snRNA-seq**) on the brain specimens and analyzed data with *cellector*, a bioinformatical approach developed to identify Mc within a predominant autochthonous (indigenous) population by analyzing variants present in the aligned reads^20^. This approach allowed for simultaneous genetic demultiplexing and high-resolution phenotyping based on transcript signatures. Finally, we sought and analyzed published snRNA-seq data of brain from healthy individuals spanning gestation to late adulthood^21–23^ including individuals of similar ages for comparison to our epilepsy patients.

## Results

### Maternal Mc prevalence in epilepsy brain with quantification by polymorphism-specific qPCR and according to brain region

In the <u>Br</u>ain <u>E</u>pilepsy (**BRE**) study, mother-child pairs were first genotyped to determine an informative polymorphism, unique to the mother, from a panel of polymorphism-specific highly sensitive TaqMan qPCR assays that we developed (mostly within HLA loci). Genotyping for 52 mother-child pairs revealed 37 pairs in which the brain could be interrogated for maternal Mc with one or more of our qPCR assays (**Table 1**). [Other mothers were homozygous or heterozygous identical (n=7), brain DNA was insufficient (n=5), or we had no qPCR assay developed for the maternal polymorphism (n=3).] Results were reported standardized to a denominator of 10^5^ as the median amount of interrogated gDNA per specimen was 94,065 gEq, [interquartile range (IQR) 70,871–111,828]. Maternal Mc prevalence was 70% with 26 of 37 patients having maternal Mc in at least one brain region. Maternal Mc was represented across regions similarly in temporal, frontal, and parietal lobes and in the hippocampus. Prevalence within regions ranged from 62% to 70%. Quantitative results were as high as 458.89 gEq/10**^5^** in a hippocampus sample. The median Mc quantitative value across samples was 2.2 [IQR 0.0–31.7] gEq/10**^5^** (**Table 1**). Thus, maternal Mc is frequently found in the brain from children with epilepsy, sometimes in substantial amounts, and broadly distributed across brain regions.

**Table 1.** Maternal microchimerism in surgically resected epilepsy brain by region. Maternal Mc was quantified in specimens from 37 informative BRE probands using qPCR assays targeting maternal-specific polymorphisms. Values are maternal Mc genome equivalents per 100,000 host genome equivalents (gEq/10⁵); 0 indicates assayed but not detected, and blank cells indicate not tested. Age is age at surgery; two probands underwent multiple resections. ‘Other’ comprises occipital tissue (BRE022 and BRE043), dysembryoplastic neuroepithelial tumor tissue (BRE025), and tissue at the edge of a ganglioglioma (BRE034). The Assay column identifies the polymorphism-specific qPCR assay; the Confirmatory assay column reports agreement for specimens assessed with more than one assay. Bold italic BRE IDs denote probands included in snRNA-seq. BRE, Brain Epilepsy; gEq, genome equivalents; Mc, microchimerism; qPCR, quantitative polymerase chain reaction; snRNA-seq, single-nucleus RNA sequencing.

| BRE ID | Temporal* | Frontal* | Parietal* | Hippocampus* | Other*± | Age^ | Assay# | Confirmatory assay# |
| --- | --- | --- | --- | --- | --- | --- | --- | --- |
| 001 |  |  |  | 316.6 |  | 1 | AT3-S | Chi0492i^^ |
| 002 | 0.0 |  |  | 0.0 |  | 0.1 | DRB1*03 |  |
| 006 | 0.0 | 9.2 |  |  |  | 9 | DRB1*15/16 |  |
| 007 | 0.0 |  | 417.5 |  |  | 0.7;5.9 | DQA1* 01 | Chi0492i^^ |
| 008 | 21.5 |  |  | 4.6 |  | 8.5 | DQA1*05 |  |
| 009 | 0.0 | 0.0 | 0.0 | 0.0 |  | 2 | DRB1*08 |  |
| 011 |  |  | 1.8 |  |  | 2.5 | DRB1*07 |  |
| 012 |  | 0.0 | 102.2 | 66.6 |  | 9 | DRB1*07 |  |
| 013 |  | 12.3 |  | 16.2 |  | 6.8 | DRB1*04 |  |
| 014 | 0.8 |  |  | 458.9 |  | 13 | DQA1*05 | AT3-L** |
| 015 |  | 53.1 | 3.8 |  |  | 3.5 | DRB1*03 | AT3-L±± |
| 017 |  | 65.8 |  |  |  | 11.2 | DQA1*05 |  |
| 020 |  |  | 0.0 | 0.0 |  | 10.1 | DRB1*01 |  |
| 021 | 6.0 |  |  | 1.1 |  | 14.1 | DRB1*04 |  |
| 022 |  |  |  |  | 144.8 | 11.8 | DQB1*03 |  |
| 023 |  |  |  | 80.2 |  | 5.5 | TNN-ins | DRB1*10^^ |
| 024 | 3.4 |  |  | 0.0 |  | 17 | DRB1*04 |  |
| 025 |  |  |  |  | 0.0 | 13 | DRB1*07 |  |
| 026 |  | 0.0 |  |  |  | 12.9 | DRB1*01 |  |
| 027 | 2.2 |  |  | 0.0 |  | 6.2 | DQA1*01 |  |
| 033 | 29.2 |  |  |  |  | 3 | DRB1*04 |  |
| 034 |  |  |  |  | 0.0 | 11.8 | DRB1*01 |  |
| 035 | 2.1 |  |  |  |  | 3.8 | DQA1*01 |  |
| 037 | 34.0 |  |  |  |  | 15 | DRB1*03 |  |
| 038 | 3.1 |  |  |  |  | 9 | DRB1*03 |  |
| 039 |  |  |  | 0.0 |  | 1.4 | DRB1*07 |  |
| 040 |  | 413.3 | 269.1 | 76.1 |  | 13.3 | DRB4*01 |  |
| 041 |  |  | 0.0 |  |  | 3.5 | DRB1*14 |  |
| 042 | 194.6 | 288.8 |  | 19.3 |  | 7.9 | DRB1*15/16 | AT3-L^^ |
| 043 |  |  |  |  | 0.0 | 8.5 | DRB1*03 |  |
| 044 | 31.7 |  |  | 25.6 |  | 9.5 | DQA1*05 |  |
| 046 | 0.0 |  |  | 0.0 |  | 15 | DRB1*07 |  |
| 047 | 0.0 |  |  |  |  | 14 | DRB1*03 |  |
| 048 | 11.4 | 2.0 |  |  |  | 17.4;18.6 | DRB1*04 |  |
| 049 | 1.6 |  |  |  |  | 17.8 | DQA1*05 |  |
| 051 | 0.0 |  |  | 1.0 |  | 10 | DRB1*01 |  |
| 054 | 5.1 |  |  |  |  | 13.9 | DQB1*02 |  |

\* genome equivalent number of microchimeric cells per 100,000 total cells (gEq/10<sup>5</sup>)
^ age at surgery (years); 2 patients underwent multiple resections
± other regions: occipital (022&043; tumor DNET(025), ganglioma edge (034)

To further substantiate the source of Mc detected as maternal in origin we conducted additional testing for brain specimens that could be studied with more than one qPCR assay (**Table 1**). Of six patients, four were concordant and in similar quantities when tested for the same region with two different assays (one of whom was concordant in each of two regions). One patient tested by two different assays was maternal Mc positive for both in one region but in differing amounts and low-level positive vs negative in another region. One patient was discordant in a hippocampus sample, positive by one assay negative by the other assay. Possible explanations for this sample include random sampling or that Mc was from a source other than maternal. The latter explanation is suggested by this mother’s history which was of ten pregnancies with multiple miscarriages and ectopic pregnancies that may have preceded the patient’s birth. Also noted were two patients who had two surgeries, both of whom were maternal-Mc-positive in the tested sample from the second surgery.

We evaluated maternal Mc results for differences according to patient sex, age at seizure onset, age at surgery, time from seizure onset to surgery, and birth order. There were 14 males and 12 females who were maternal Mc positive versus 5 males and 6 females who were maternal Mc-negative. Age at seizure onset was similar in maternal Mc positive and negative groups, respectively, 0 to 14 years versus 0 to 16 years. Age at surgery was distributed similarly in maternal Mc positive and negative groups, respectively, 0 (birth) to 14 years versus 28 days to 18 years. Time from seizure onset to surgery did not significantly differ in the two groups: 6 months to 17.8 years (median 5 years) in the former against 28 days to 7 years in the latter (median 2 years). Birth order, however, differed in the two groups. Among maternal Mc positive patients, 14 were first born and 12 second or later born contrasting with maternal Mc negative patients with just one first born and 10 second or later born. This trend suggests possible displacement/replacement by different sources of Mc “grafts” as has been demonstrated in a murine model^24^ and theorized as an aspect of evolutionary conflicts^25^ including sibling rivalry^26^.

### Maternal Mc and Mc agnostic to source by single-nucleus RNA sequencing in epilepsy and healthy brain

Next we conducted massively parallel single cell transcriptomic sequencing to identify microchimeric cells through transcriptome wide variant analysis. Of the 37 patients with maternal qPCR results, 10 patients had sufficient excess frozen brain tissue for transcriptome profiling. We performed single-nucleus RNA droplet capture of 11 brain tissues (9 patients) using 10x Genomics 3’ single-cell chemistry and of 4 brain tissues (3 patients already tested in 3’ and an additional 10th) using 10x Genomics 5’ GEM-X single cell chemistry. Rationale for the latter chemistry involved the assumed better gene-per-cell sensitivity and cell recovery compared to older chemistries and capability to double the number of brain cells assessed^27^. We captured a median of 12,473 high quality nuclei [IQR 9,929–14,760] in the 3’ capture and 27,671.5 [IQR 19,304.5–36,339.5] in the 5’ capture, after thorough nuclei quality control including two consecutive passages through scrubletR^28^ to remove ‘multiplet’ artifacts (see methods).

To contextualize the epilepsy findings, we analyzed 41 publicly available healthy and cognitively normal human brain specimens generated using 10x Genomics 3′ snRNA-seq. Herring et al.^21^ profiled postmortem brain specimens from 29 individuals without reported neuropathology or neurodevelopmental disorders spanning 22 weeks of gestation to 40 years of age and including individuals whose ages overlapped with those in our epilepsy cohort. To extend the reference population into later adulthood, we also analyzed nine cognitively normal specimens from the Alzheimer’s disease study by Lau et al.^22^ and three from the multiple sclerosis study by Feng et al.^23^ Raw sequencing data from all three studies were reprocessed using the same analytical pipeline applied to our epilepsy datasets to enable uniform comparison. Overall, the number of nuclei passing quality control totaled 337,964 and had a median of 7,384 [IQR 5,437–10,208] per sample, in line with the originally published data^21–23^.

To identify Mc in the cell-barcoded transcriptomic data, we used *cellector*, a genetic demultiplexing algorithm capable of distinguishing allogeneic cells at remarkably low frequencies by analyzing expressed genetic variants present in the aligned reads^20^. *Cellector* uses beta-binomial distribution to model the majority genotype, scores the log likelihood that a cell’s observed variant pattern matches these under a fitted model, and then iteratively applies anomaly detection (interquartile range outlier-based) to identify minority/foreign-genotype cells within a majority population.

Mc was identified in both the epilepsy and the healthy brain datasets. Overall, Mc was detected in 64% (36/56) of the captured single-nucleus from specimens across all datasets by *cellector* (**Fig. 1A–1D**) Counts were as high as 65 and 46 microchimeric nuclei in the hippocampus and temporal lobe specimens of an epilepsy patient, BRE044 (**Fig. 1D**). Per-locus likelihood summaries indicated that Mc assignments were supported by contributions from multiple expressed variants across specimens (**Suppl. Fig. S1**). As an additional approach, we used *souporcell*, a well-established genetic demultiplexing algorithm designed to distinguish allogeneic cells in proportionate/balanced mixtures, using variants present in the aligned reads^29^. In this epilepsy patient’s specimens, nuclei forming a distinct component in *souporcell’*s sparse mixture model output were predominantly classified as Mc by *cellector*, demonstrating concordance between the two variant-based classification approaches (**Fig. 1E**). Comparable separation was not observed in specimens with lower Mc counts, consistent with the limited ability of souporcell to resolve highly imbalanced genotype mixtures. Because this epilepsy patient was male, we further examined expression of the female-associated long noncoding RNA *XIST*. *XIST* was detected in most candidate microchimeric nuclei but rarely in host nuclei (**Fig. 1F**), supporting a female source and, in conjunction with the maternal-specific qPCR result and the patient’s young age (9 y/o), a likely maternal origin. *XIST* expression was not detected in microchimeric nuclei from the remaining male probands, likely reflecting sparse transcript detection in snRNA-seq data.

**Figure 1.**
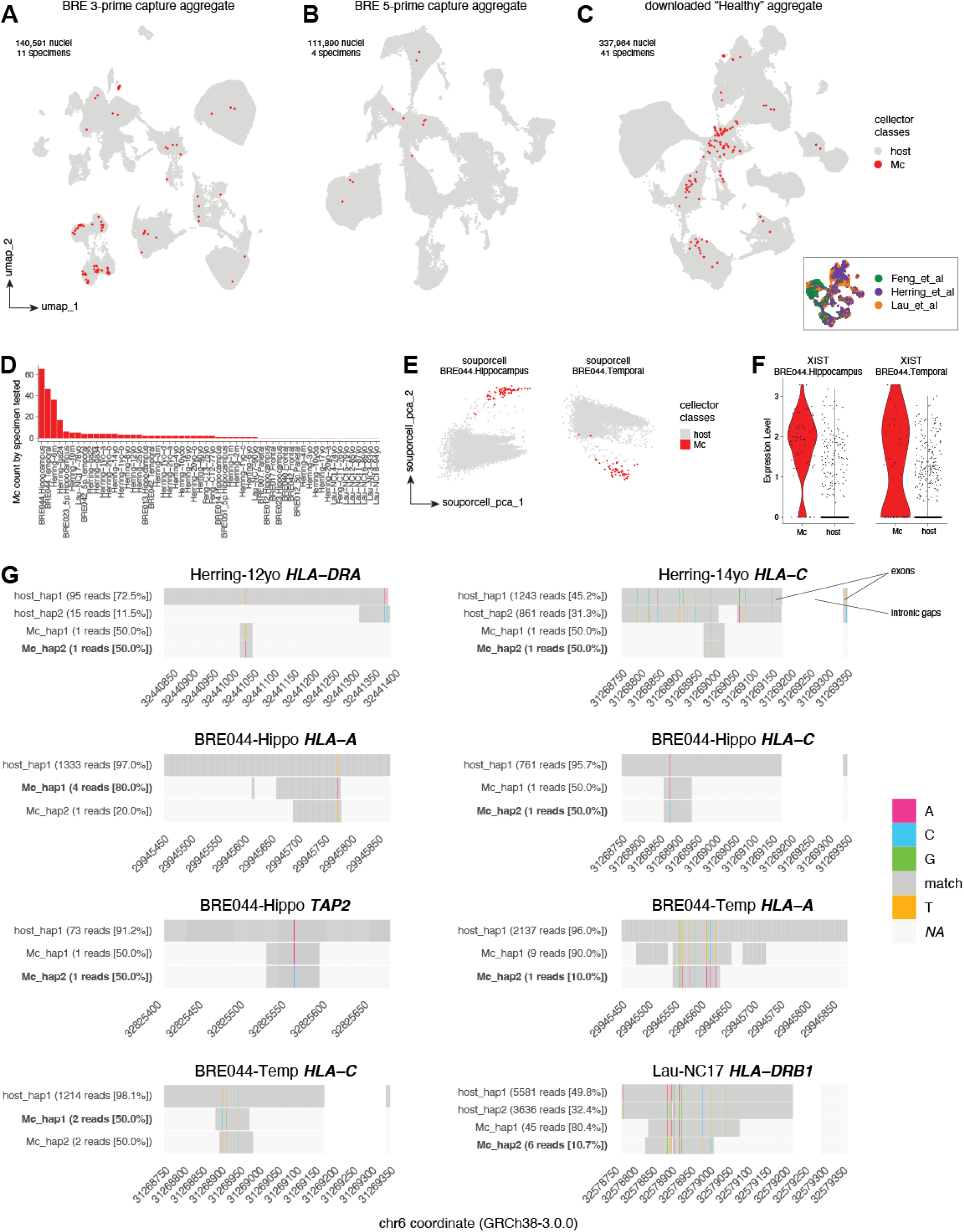
Variant-based identification (cellector) and genetic characterization of microchimeric nuclei in human brain. **A-C,** Uniform manifold approximation and projection (UMAP) embeddings of aggregated single-nucleus RNA-sequencing datasets from Brain Epilepsy (BRE) specimens captured using 10x Genomics 3′ chemistry (**A**; 140,591 nuclei from 11 specimens and 9 probands), BRE specimens captured using 5′ chemistry (**B**; 111,890 nuclei from 4 specimens and 4 probands), and publicly available neurotypical healthy specimens captured using 3′ chemistry (**C**; 337,964 nuclei from 41 “healthy” probands). Nuclei are colored by cellector classification as majority-genotype host nuclei (gray) or candidate microchimeric nuclei (red). The inset in **C** shows the control aggregate colored by source study: Feng et al. (green), Herring et al. (purple), and Lau et al. (orange). **D,** Number of microchimeric nuclei identified by cellector in each of the 56 single-nucleus captures, ordered by observed Mc count. **E,** Principal-component representation of the log-normalized souporcell sparse-mixture output for hippocampal and temporal-lobe specimens from proband BRE044. Points are colored by the corresponding cellector classification, showing concordance between the two variant-based classification approaches. **F,** Distribution of normalized expression of the female-associated long noncoding RNA *XIST* among Mc and host nuclei in the BRE044 hippocampal and temporal-lobe specimens. Points represent individual nuclei. **G,** Representative read-backed haplotype reconstructions at informative HLA and HLA-associated loci in epilepsy and control probands. Rows show supported haplotypes reconstructed independently from host and Mc reads; parentheses give the number and percentage of reads assigned to each displayed haplotype. Gray tiles indicate matching bases at mutually covered positions, colored tiles indicate discordant nucleotides (A, magenta; C, cyan; G, green; T, orange), and white indicates unavailable coverage. Genomic positions refer to chromosome 6 in GRCh38-3.0.0. HLA, human leukocyte antigen; Mc, microchimeric; PCA, principal component analysis; UMAP, uniform manifold approximation and projection.

To evaluate genetic relatedness between microchimeric and host nuclei, we performed read-backed phasing of expressed variants at HLA loci. This choice was made because HLA genes are located in one of the most variant-dense and polymorphic stretches of the human genome^30^, making haplotype phasing possible despite the limitation of working on short-read sequencing and transcriptomic data. However, because of the significant HLA transcript drop out (**Suppl. Fig. S2**), which is thought to be an artifact of nuclei preparation and capture^31^, sufficient coverage for HLA variant phasing analysis was available for 9 of 51 epilepsy and healthy probands (see methods). At informative loci, Mc reads carried both one shared and one non-shared haplotype with the host (**Fig. 1G**) and, in some instances, only one shared haplotype or only one non-shared haplotype with the host were observed (**Suppl. Fig. S3A–B**). A few other loci were uninformative because of limited variant availability in the covered positions (**Suppl. Fig. S3C**). No unexpected haplotype configuration was observed (e.g., Mc population containing two non-shared haplotypes), a pattern consistent with first-degree genetic relatedness.

Single cell transcriptomic data both confirm and extend the observation that Mc is common in epilepsy brain as assessed by polymorphism-specific qPCR. Transcriptomic data analysis of healthy brain further indicates Mc is common in healthy human brain. Additionally, while limited transcript coverage and absent parental genotypes preclude definitive Mc origin assignment, concordance between variant-based classifications, *XIST* expression in a male patient, and HLA-phased variant haplotype sharing in informative probands collectively support genetic relatedness of Mc and host, consistent with frequent maternal origin as observed in epilepsy.

### Heterogeneous and wide-ranging transcriptomic profiles of Mc in epilepsy and healthy brain

To characterize the cellular identities and transcriptional profiles of microchimeric nuclei, we annotated all nuclei with viewmastR^32^ using the well-annotated Allen Brain Map Human M1 Cortex Atlas^33^ as a reference (**Fig. 2A**; Methods). Across cohort-level aggregates, viewmastR recovered the major neuronal, glial, and vascular cell classes. In epilepsy specimens, annotated cell-type proportions varied across brain regions and replicate captures, including paired brain regions from the same proband and paired 3’ and 5’ libraries from the same region and proband, potentially reflecting regional biology, sampling variability, and capture chemistry effects (**Suppl. Fig S4A**). In the healthy datasets, the broad cell-class assignments were consistent with those reported in the source publications. Particularly within the Herring series^21^ relative oligodendrocyte representation generally increased from infancy to adulthood^34^ (**Suppl. Fig. S4B**).

**Figure 2.**
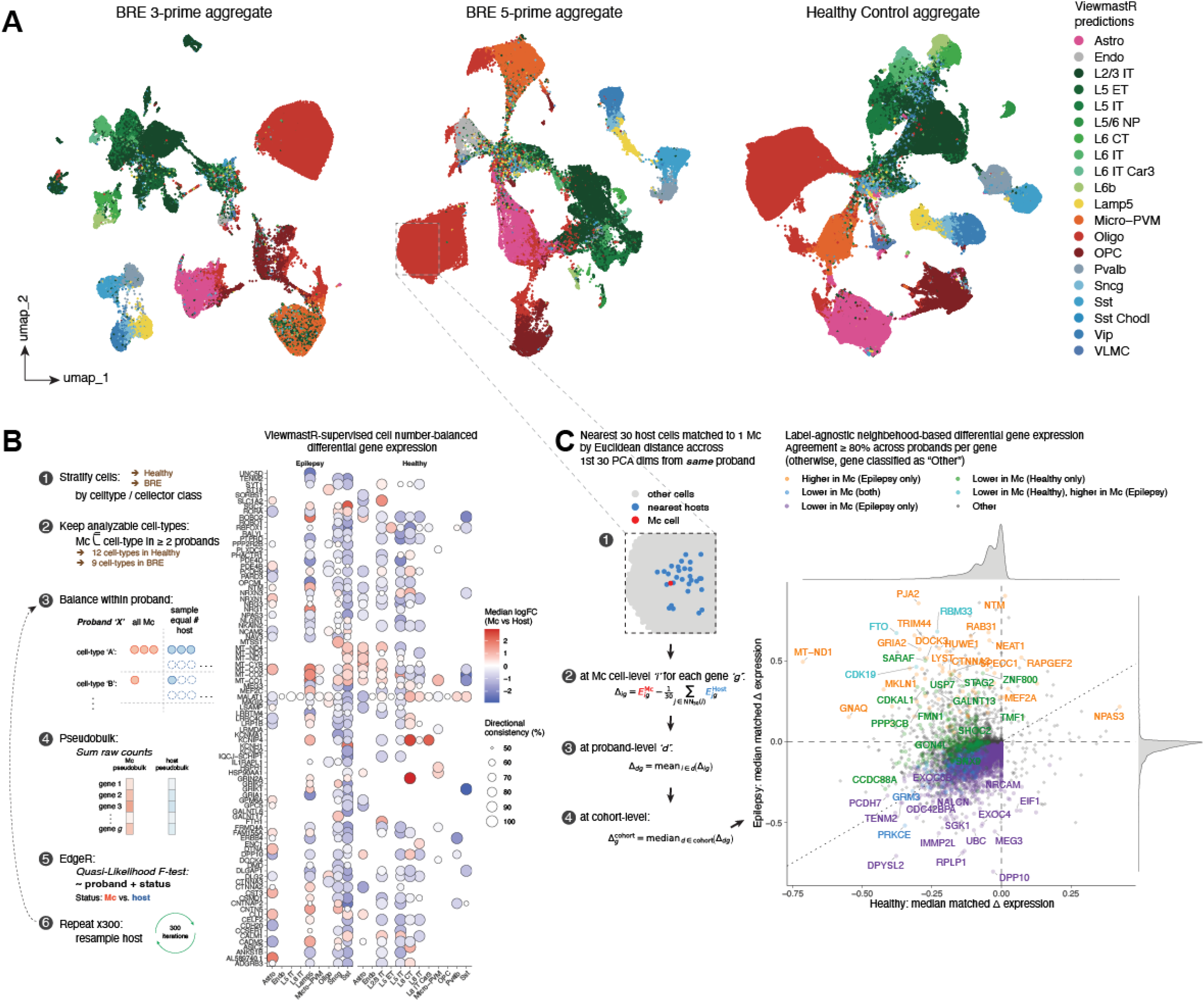
Cell-type identities and transcriptional differences of microchimeric nuclei. **A,** UMAP embeddings of the BRE 3′, BRE 5′, and healthy aggregates colored by viewmastR-predicted cell type. Cell identities were assigned using the Allen Brain Map Human M1 Cortex single-nucleus atlas as the reference. **B,** Cell-type-supervised, cell-number-balanced Mc-host expression analysis. Nuclei were stratified by viewmastR identity and cellector class, and the analysis was restricted to cell types containing candidate microchimeric nuclei in at least two independent probands, yielding 9 analyzable cell types in BRE and 12 in the healthy cohort. Within each informative proband, and cell type, all microchimeric nuclei were retained and an equal number of host nuclei was randomly sampled. Raw counts were summed to form paired Mc and host pseudobulks and analyzed with an edgeR quasi-likelihood model containing proband and Mc status. Host sampling was repeated 300 times. For each displayed gene-cell-type combination, bubble color indicates the median log2 fold-change across iterations (red, higher in Mc; blue, lower in Mc), and bubble diameter indicates the percentage of iterations with a concordant direction of effect. **C,** Label-agnostic matched-neighbor expression analysis. Each Mc nucleus was compared with its 30 nearest host nuclei from the same proband using Euclidean distance across the first 30 expression-derived principal components. For each gene, normalized expression in the microchimeric nucleus minus the mean expression of its matched host neighbors was averaged within proband and summarized by the median across probands. The scatterplot compares cohort-median differences in healthy (x axis) and epilepsy (BRE) specimens (y axis); marginal density plots show the corresponding distributions. Horizontal and vertical dashed lines indicate zero, and the diagonal dashed line indicates equal cohort-median differences. Colors denote directional concordance in at least 80% of Mc-positive probands (5 of 6 Mc+ BRE probands and 23 of 28 Mc+ healthy probands): orange, higher in Mc in BRE only; blue, lower in Mc in both cohorts; purple, lower in Mc in BRE only; green, lower in Mc in healthy only; cyan, lower in Mc in healthy and higher in Mc in BRE; and gray, all other patterns. Color categories indicate direction rather than statistical significance. Astro, astrocyte; BRE, Brain Epilepsy; Chodl, chondrolectin-expressing; CT, corticothalamic; Endo, endothelial; ET, extratelencephalic; IT, intratelencephalic; L, cortical layer; Lamp5, LAMP5-expressing; logFC, log2 fold-change; Micro-PVM, microglia/perivascular macrophage; NP, near-projecting; Oligo, oligodendrocyte; OPC, oligodendrocyte precursor cell; Pvalb, Parvalbumin-positive; PCA, principal component analysis; Sst, Somatostatin-expressing; Vip, vasoactive intestinal polypeptide-expressing; VLMC, vascular and leptomeningeal cell.

We next tested Mc–host expression differences within annotated cell types, analyzing the epilepsy and healthy cohorts separately. Cell-number-balanced, proband-level pseudobulk counts were evaluated using an edgeR quasi-likelihood model^35,36^ that accounted for pairing by proband (**Fig. 2B**). The analysis was restricted to cell types containing microchimeric nuclei in at least two independent probands, yielding nine analyzable cell types in the epilepsy cohort and 12 in the healthy cohort. Among the genes highlighted by this analysis in both cohorts, negative Mc–host logFC (log2 fold-change) estimates frequently involved genes associated with synaptic adhesion and circuit organization, including *PTPRD*, *NRXN1/3*, *CNTNAP2*, *TENM2*, *LRRTM4* and *ROBO1/2*, as well as genes associated with ion-channel function, glutamate signaling, and neuronal RNA regulation, including *DPP10*, *GRIA1*, *GRIK1/2*, and *RBFOX1*^37–39^. We observe fewer genes associated with a positive Mc-host logFC, but these include genes associated with ion-channel function, e.g., *KCNIP4* and glutamate receptors, e.g., *GRIN2A* implicated in a broad spectrum of neurodevelopmental/neurologic phenotypes, including epilepsy^40^.

Because the cell-type-stratified analysis excluded cell types represented by microchimeric nuclei in only one proband (biological replication was at the proband level), we complemented it with a label-agnostic, within-proband, matched-neighbor, across full Mc population analysis. Each microchimeric nucleus was compared with its 30 nearest neighboring host nuclei from the same proband using Euclidean distance across the first 30 gene expression-derived principal components. For each gene, the Mc-minus-host normalized-expression difference was calculated for each microchimeric nucleus, averaged within each proband, and summarized by the median across probands (**Fig. 2C**). Directional categories required concordant differences in at least 80% of probands (i.e., 5 of the 6 Mc-positive epilepsy and 23 of the 28 Mc-positive healthy probands).

Among the 11,582 genes analyzed, 84.7% in the epilepsy cohort and 78.4% in the healthy cohort had negative cohort-median Mc–host differences, whereas 7.1% and 0.8%, respectively, had positive differences. The remaining genes had cohort-median differences of zero. Thus, lower normalized expression values predominated in both cohorts, while positive differences were descriptively more frequent in the epilepsy cohort. Marginal densities were concentrated near zero, indicating that most cohort-median differences were small. Distance from the origin reflected effect magnitude, whereas color denoted directional concordance across at least 80% of probands. From these findings, we conclude that as a general trend, microchimeric nuclei have fewer transcripts for most genes than host nuclei. However, within the context of epilepsy this state of reduced transcription in microchimeric nuclei is less absolute.

In the epilepsy cohort, genes showing concordant positive differences included neuronal/synaptic genes such as *NTM*, *GRIA2* and *DOCK3*, and RNA-regulatory and proteostatic genes including *FTO*, *HNRNPA2B1*, *NEAT1*, *HUWE1* and *PSME4*^41–43^. Notably, the concomitant increase in *GRIA2* and decrease in *DPP10* implicate pathways controlling glutamatergic transmission and membrane excitability, both of which are strongly coupled to epileptic network activity^44,45^. Additionally, *MEF2A*, another gene showing concordant positive difference in epilepsy cohort, is a key transcription factor that controls expression of numerous genes involved in synaptic and neuronal function. Notably, therapeutic downregulation of *MEF2A* may ameliorate epileptic seizures^46,47^. In the healthy cohort, genes showing concordant lower expression in microchimeric nuclei included *CCDC88A*, *PPP3CB*, *CDKAL1*, *FMN1*, and *PRKCE*, which have reported roles in cytoskeletal organization, neuronal signaling, and cellular architecture^41,48,49^.

We next applied Hallmark gene-set enrichment analysis (**GSEA**) to the matched-neighbor expression differences calculated separately for the epilepsy and healthy cohorts. Although no Hallmark pathways reached conventional significance (false discovery rate < 0.05), GSEA revealed suggestive and directionally divergent enrichment of Mc between epilepsy and healthy brain, most notably for Hedgehog signaling and mitotic spindle (**Suppl. Fig. S5**), but insufficient to infer disease-associated differences in Mc transcriptional programs.

Overall, both cell-type-supervised and label-agnostic analysis strategies supported a Mc associated alteration in neuronal structural and functional state, but not necessarily a uniform disease-specific transcriptional program. These findings suggest either altered regulation of transcription and/or transcript stability as a potential feature of microchimeric nuclei–an idea that will require further biological validation.

### Mc enrichment in neuronal subtypes and microglia, with quantitative age-dependent variation over the human lifespan

We next quantified cellector-identified Mc as the number of candidate microchimeric nuclei by total nuclei number. Across 15 epilepsy captures from 10 probands, median Mc levels were 5.9 [IQR 0.0–15.4] nuclei/10**^5^** (**Suppl. Fig. S6A**). Although the cellector and maternal-specific qPCR estimates were broadly similar in scale, a descriptive specimen-level comparison did not show a monotonic association. One explanation could be that qPCR analyzed approximately 10-fold more cell equivalents than snRNA-seq, providing greater quantitation accuracy with fewer cells tested in snRNA-seq data translating into a wider 95% confidence interval (95%CI) based on the Wilson score without continuity correction^19^ as we observed (**Suppl. Fig. S6B**). Data from the DNA-based and the RNA-based approaches is also expected to differ based on differential gene expression according to developmental stage and age. Differences may also reflect transcript detection, qPCR efficiency, tissue sampling, and the distinction between source-agnostic cellector classification and maternal-specific qPCR measurement.

Microchimeric nuclei were assigned diverse neuronal, glial, immune, and vascular cell-type identities (**Suppl. Fig. S6A**), and their distribution did not consistently parallel the overall composition of host population (**Suppl. Fig. S4**). For proband-level analyses, we pooled multiple brain-region and capture-chemistry datasets from the same epilepsy proband by summing microchimeric and total-nucleus counts before calculating Mc frequencies (**Fig. 3A, 3B**). The resulting median among epilepsy probands was 7.3 [IQR 0.0–16.2] nuclei/10**^5^**. Among 41 healthy probands the corresponding median was 24.4 [IQR 0.0–45.3] nuclei/10**^5^**.

**Figure 3.**
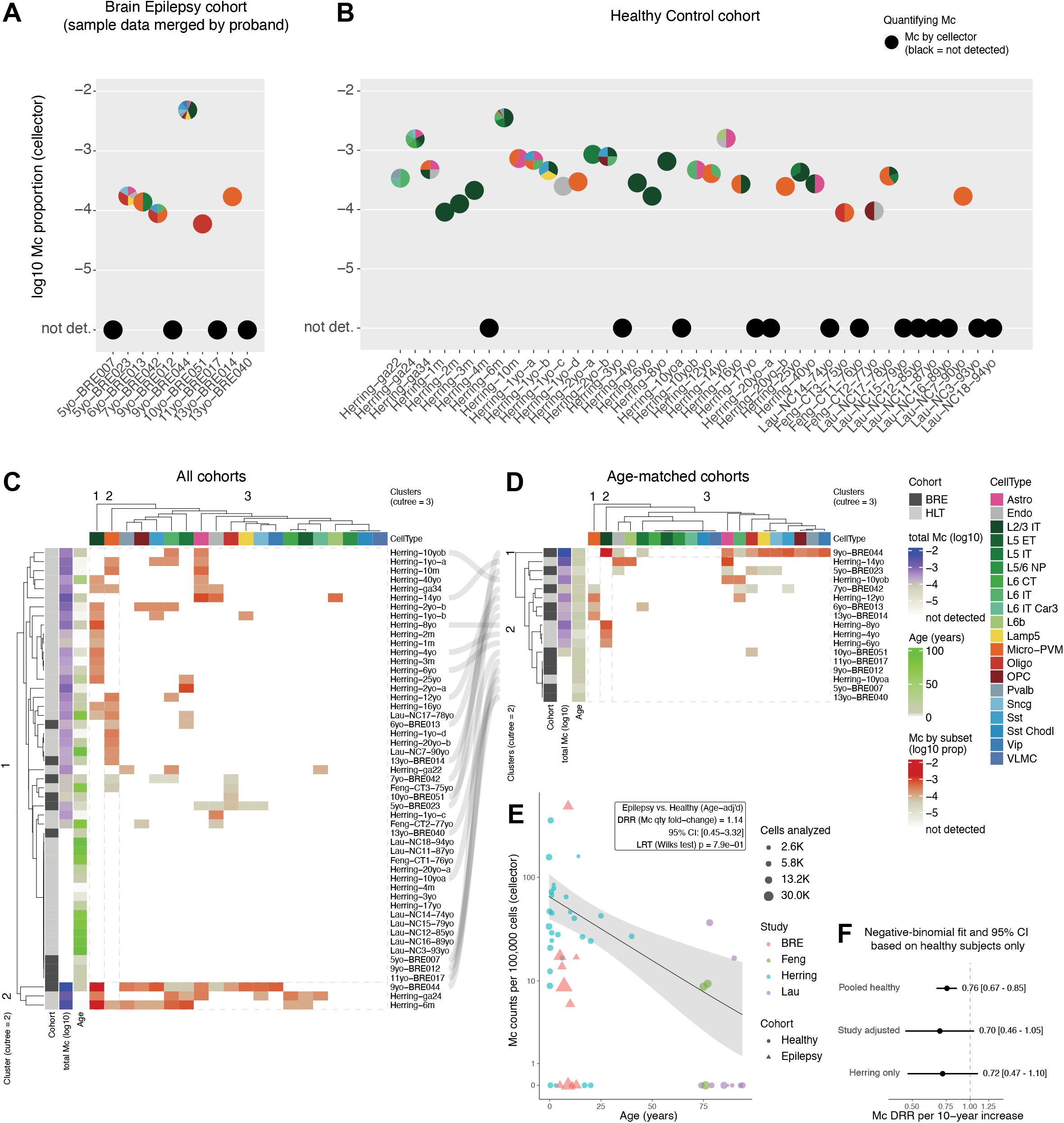
Cell-type distribution and age-associated quantification of microchimeric nuclei. **A, B,** Proband-level candidate Mc frequencies and cell-type distributions in the displayed BRE epilepsy probands (**A**) and 41 healthy probands from the Herring, Feng, and Lau studies (**B**). The vertical position of each pie chart represents log10 (microchimeric nuclei/total nuclei), and pie sectors show the viewmastR-predicted cell-type composition of microchimeric nuclei. Black points at ‘not detected’ denote probands in whom cellector identified no microchimeric nuclei. **C, D,** Exploratory complete-linkage hierarchical clustering of cell-type-specific Mc frequencies in all probands (**C**) and after restricting healthy probands to those within one year of the BRE age range (**D**). Heatmap cells show log10 (microchimeric nuclei assigned to the indicated cell type/total nuclei examined); white denotes no detected nuclei of that Mc-cell-type combination. Euclidean distances were used for row and column clustering, and the dendrograms were cut at k = 2 for probands and k = 3 for cell types. The column cut separated L2/3 intratelencephalic neurons and Micro-PVM as individual clusters from the remaining cell types. Side annotations indicate cohort, overall log10 Mc frequency, and age, and the top annotation identifies cell type. Gray ribbons connect probands retained from the complete analysis in **C** to their positions in the age-restricted analysis in **D**. **E,** Mc counts per 100,000 nuclei plotted against proband age. Color denotes source study, shape denotes cohort, and point area represents the total number of nuclei examined. The black line and gray band show the pooled-control negative-binomial age fit and its 95% confidence interval. Models included log10 total nuclei examined as an offset. In the exploratory age-adjusted epilepsy-versus-healthy comparison, the Mc detection-rate ratio (DRR) was 1.14 (95% CI, 0.45-3.32; likelihood-ratio test, P = 0.79). **F,** Control-only negative-binomial estimates of the Mc DRR per 10-year increase in age. Points and horizontal lines show estimates and 95% confidence intervals for pooled controls. The vertical dashed line marks the null value, DRR = 1. Astro, astrocyte; BRE, Brain Epilepsy; Chodl, chondrolectin-expressing; CI, confidence interval; CT, corticothalamic; DRR, detection-rate ratio; Endo, endothelial; ET, extratelencephalic; IT, intratelencephalic; L, cortical layer; Lamp5, LAMP5-expressing; LRT, likelihood-ratio test; Mc, microchimerism; Micro-PVM, microglia/perivascular macrophage; NP, near-projecting; Oligo, oligodendrocyte; OPC, oligodendrocyte precursor cell; Pvalb, Parvalbumin-positive; Sst, Somatostatin-expressing; Vip, vasoactive intestinal polypeptide-expressing; VLMC, vascular and leptomeningeal cell.

Visual inspection of the proband-level profiles suggested that Mc identities were dominated by microglial annotations in some probands and neuronal annotations in others (**Fig. 3A, 3B**). This aligns with other studies showing Mc of maternal origin in microglia in experimental models^15^, donor-derived Mc in microglia after allogeneic hematopoietic cell transplantation^17^, fetal cells in mouse models acquiring neuron and other brain cell phenotypes^13^, and donor-derived neurons in the cortex and hippocampus following bone marrow transplantation in humans^50^. These data raise the question whether Mc preferentially acquires certain cell fates according to the timing and location of developmental entry or disease state.

To formalize the process, we performed an exploratory analysis of recurrent cell-type-specific Mc patterns. For epilepsy probands represented by multiple regions or capture chemistries, datasets were pooled as described above. Complete-linkage hierarchical clustering using Euclidean distance was applied to the cell-type-specific Mc frequencies, with the column and row dendrograms cut at (k=3) and (k=2), respectively, to summarize the coarse structure of the sparse data. The column-wise cut separated L2/3 intratelencephalic (L2/3 IT) and microglial/perivascular macrophage (Micro-PVM) annotations from the remaining cell types (**Fig. 3C**). The row-wise cut largely separated probands with higher and lower total Mc frequencies. L2/3 IT neuron–dominant profiles occurred primarily among younger healthy probands, whereas Micro-PVM-dominant profiles occurred among older probands from both cohorts. The epilepsy proband with the highest estimated Mc frequency clustered with two young control probands (one sampled at 24 gestational weeks and one at 6 months of age) who also had relatively high Mc frequencies.

Because the healthy cohort spanned a substantially broader age range than the epilepsy cohort, we repeated the clustering after restricting healthy probands to within one year of the epilepsy age range (**Fig. 3D**). The age-restricted analysis displayed a similar coarse separation of L2/3 IT, Micro-PVM, and the remaining annotations, while the epilepsy proband with the highest Mc frequency formed the sole member of the higher-frequency row cluster. L2/3 IT neuron–dominant profiles occurred mainly among healthy probands, whereas Micro-PVM-dominant profiles occurred in both cohorts. These exploratory analyses identify L2/3 IT and Micro-PVM annotations as conspicuous components of Mc profiles, although establishment of definite enrichment, developmental fate, and preferential acquisition of these identities requires future validations.

To test the effect of age, we modeled Mc counts with respect to age in a negative binomial regression^51^ with total number of cells examined included as an offset (log-total counts). The statistical model showed that Mc rates declined with age among 41 healthy subjects (detection rate ratios [DRR] per decade, 0.76; 95%CI, 0.67–0.85; likelihood-ratio test [LRT], *p* = 4.7e-5; **Fig. 3E, 3F**). The association remained similar in magnitude after adjustment for study (DRR, 0.70; *p* = 0.091) and within the Herring cohort alone (DRR, 0.72; LRT, *p* = 0.134), although with reduced precision (**Fig. 3F**). Given this age association, we next compared Mc rates between epilepsy and healthy brains while accounting for age and found no evidence of a difference (epilepsy vs. healthy DRR, 1.14; 95%CI, 0.45–3.32; LRT, *p* = 0.79; **Fig. 3E**). Because all epilepsy samples originated from the disease cohort, however, disease status remained confounded with study.

Considering these results and analyses overall, it is apparent that Mc is prevalent in the human brain, manifests in diverse cell types, preferentially in microglia and a subtype of neurons, can be present in substantial quantities, persists for decades, likely throughout a human lifetime, and declines with age.

## Discussion

In this study we report that Mc is a prevalent, diverse, and long-lived cellular feature of the human brain. Using a panel of polymorphism-specific qPCR assays to target maternal-specific polymorphisms we found maternal Mc in multiple brain regions in a majority of epilepsy brain specimens, often at high levels. Maternal Mc is acquired early in life, reported in a normal human fetus at 13 weeks gestation^52^, in mice at day 12.5 gestation^14^, and may also be acquired postnatally from breastmilk^53^. Our study population for maternal Mc quantification by polymorphism-specific qPCR consisted of pediatric epilepsy patients with an age range from birth to 16 years at seizure onset. Although pathogenic pathways leading to epilepsy are likely multiple, some share biological links with neurodevelopmental disorders and multiple studies have reported maternal Mc effects on development^54^. Moreover, a recent murine study demonstrated maternal Mc can impact neurodevelopment, brain homeostasis, synaptic pruning, and behavioral abilities^15^.

Conducting single-nucleus sequencing, we found that maternal cells were present as neurons, multiple neuronal subtypes, oligodendrocytes, astrocytes, microglia, and endothelial cells. Mc enrichment within microglial/perivascular macrophage-like cells and L2/3 intratelencephalic neurons raises the possibility that developmental windows and tissue niches shape fates of maternal cells in the brain. Microglia enter the brain from extraembryonic structures early in gestation^55^ whereas L2/3 IT neurons are late-born upper-layer cortical projection neurons generated during later waves of corticogenesis^56^. Interestingly, Mc was not detected among L5/6 near-projecting neurons, a deeper-layer excitatory population associated with earlier cortical developmental programs^57,58^, nor among Sst Chodl neurons or VLMCs, which follow distinct embryonic somatostatin projection-neuron^59^ and meningeal mesenchymal^60^ developmental programs, respectively. Thus one possibility is that maternal progenitor cells (or maternal-fetal fusion events), become incorporated during a developmental window favoring upper-layer intratelencephalic neuronal identity^61^. Studies of mice and marmosets suggest chimeric cells preferentially acquire myeloid lineage fates in the brain^15,62^ and, in humans, microglia and/or immune cell infiltrates have been reported in transplantation studies^17^. Our data, while in agreement with these observations, support multipotency of cell fates as neuroectodermal (neurons, glia), endothelial, and myeloid (microglia) cells were all identified indicating naturally acquired Mc in human brain is more prevalent and diverse than had been anticipated. The abundance of Mc in some brain regions and absence in others raises the possibility that developmental timing and lineage-specific niche accessibility constrain the cell fates available to maternal cells entering the fetal brain; a hypothesis to be explored in more detail in a future study.

Results showed overall strong transcriptional resemblance of maternal Mc to, and co-embedded with, the autochthnous cell population. Complementary cell-type-supervised and label-agnostic differential gene expression analyses identified recurrent differences in genes associated with neuronal morphogenesis, synaptic organization, membrane excitability, and RNA and protein homeostasis. These differences varied among cell types and between epilepsy and healthy brains, while the genome-wide predominance of negative expression differences could partly reflect lower molecular recovery from microchimeric nuclei in some datasets (**Suppl. Fig S7**). Thus, our findings did not support a uniform or disease-specific Mc transcriptional program but instead suggested that brain Mc comprised transcriptionally heterogeneous cells with molecular features that vary according to cell identity and tissue context. However, we did notice that compared to host cells, microchimeric cells had a generally dampened amount of gene transcripts. A common hypothesis in the field of Mc research is that maternal Mc may have a higher somatic burden due to their age and thus confer some disadvantages to their progeny, while fetal Mc may exert rejuvenating effects due to their younger age. There is some evidence supporting this in parabiosis studies between young and old animals that observed cognitive decline in the former and improvement in the latter^63,64^. Thus, repression of transcripts in microchimeric cells, combined with the observed decline in their number in aging brains, may represent endogenous mechanisms promoting healthy brain development and aging. That we observed increases in numerous gene transcripts in microchimeric cells of epileptic brains suggests that dysregulation of this process may contribute to a disease state, but further studies are needed to carefully elucidate the role and influence of Mc on brain development and disease.

While the functional significance of maternal Mc in epilepsy brain is presently unknown, one possibility is that maternal Mc migrates to sites of brain injury in the fetus or neonate, analogous to what has been shown for fetal microchimeric cells in the murine maternal brain. Fetal microchimeric cells were found to traffic to excitotoxic lesions in postpartum maternal brain in mice^65,66^. On the other hand, other studies point to a wider range of possibilities. In addition to shaping neurodevelopment, maternal Mc has been shown to partially rescue interleukin 2 deficiency, promote neonatal immunity against early life infections, modulate malarial disease severity in children, enforce reproductive fitness in females, and instruct offspring immune cells in response to self antigens^1,15,67–72^.

The biology of an individual life and the Mc harbored is dynamic, subject to change as time elapses and according to location and microenvironment among other variables^1,61^. This dynamic exists as a relationship and some mother-child HLA-relationships have been reported to increase maternal Mc in the child^73^. Additionally, females who become pregnant acquire Mc from their pregnancies. Moreover, maternal Mc replacement by Mc of fetal origin (or displacement) has been well demonstrated in mice^24^. Adding another level of complexity, occasionally Mc can be acquired from an older sibling because a mother harbors Mc from previously born children, or an earlier maternal pregnancy loss, twin, or rarely a maternal grandmother^1^. These additional sources of Mc are supported by the finding of male cells in female newborns and fetuses^74^. In our study maternal Mc was detected more often in first born vs. later born children; maternal Mc replacement by sibling Mc, although not yet described, offers a potential explanation for this observation.

Using *cellector* to interrogate public snRNA-seq datasets we further report that brain-resident Mc is detectable in healthy individuals from fetal life to advanced age. Young brains harbored Mc with neuronal features more often and older brains Mc with microglial characteristics. The high prevalence of Mc in healthy brains from the publicly available snRNA-seq datasets^21–23^ support a key finding from the epilepsy studies: that naturally acquired Mc is prevalent in the human brain and across a multitude of cell fates. This indicates the range of Mc fates we observed in epilepsy brains is not explained by disease or neurodevelopmental pathology. Further investigation of all stages of the human lifespan are needed to elucidate overall trends in Mc cell fates and assess potential differences in Mc prevalence and/or fate due to aging and/or disease status.

There are limitations to our study. We could not test brain bank tissues for maternal Mc due to lack of maternal genotype information which, even if available, could differ for post-mortem tissue from our surgically resected tissue, though other insights have been gained from interrogating post-mortem brain^75^. Thus, our maternal-specific qPCR and snRNA-seq data derives from brains affected by epilepsy. A limitation of our qPCR dataset is we were unable to test patients for whom the mother was homozygous or heterozygous identical with the patient. Studies suggest maternal Mc is greater in child-mother pairs with these HLA-relationships^73,76^ therefore maternal Mc may be underestimated. A limitation of analyses from publicly available snRNA-seq is that Mc origin is unknown as *cellector* is agnostic to Mc origin. Some technical limitations include the sparsity of single-nucleus transcriptomic data (generally so in short-read single-cell omics), and the lower quantitative accuracy of genetic demultiplexing at low cell frequencies compared to targeted qPCR. We have seen this exemplified by a clear association of the female-specific *XIST* signal only in the individual with the highest Mc content by *cellector*. Thus, depending on a single feature to detect a rare cell population is less robust.

At the same time there are strengths to the data and complementary approaches used of polymorphism-specific qPCR and single-nucleus transcriptomics. Data and analyses provide snRNA-seq profiling of Mc for the first time in the human brain. Our approach provides compelling evidence that using next generation techniques and novel bioinformatics approaches can identify rare allogeneic cells with exceptional detail into the transcriptional profile and identity of these cells. snRNA-seq data was recently reported for sibling chimerism in marmosets, however this study required whole exome genome sequencing to distinguish chimerism, and the unique biology of marmoset pregnancy results in *macro*chimerism^62^. An advantage of the *cellector* approach reported here is that it does not require prior knowledge of polymorphisms and/or other genome variants and is therefore broadly applicable to snRNA-seq analysis. In our study successful application of this approach to publicly available snRNA-seq data identified Mc in post-mortem tissue from individuals ranging from gestational ages to 93 years of age. The findings in the publicly available data from healthy brains confirm our initial key findings in epilepsy patients indicating Mc is present in the brain throughout life with a myriad of cell fates.

Our findings, considered together with other studies, has broad implications for a number of diseases. Of special interest is autism as there is strong evidence for maternal-fetal interaction in autism which, to our knowledge, is unstudied for Mc. Maternal autoantibodies have been associated with autism in a number of studies^77,78^. Moreover, an experimental murine model recently directly demonstrated that Mc can be the origin of disease-specific autoantibodies^79^. Of interest, female cells in male brain, presumed to be maternal, were described in a small number of patients with the autoimmune disease multiple sclerosis, and in a few male controls^10^. Recent appreciation of autoantibodies in numerous neurological diseases^80^ supports a rationale for further investigation of Mc in other neurological diseases. Also of interest for future study is potential consequences of a Mc “graft-vs.-graft” effect in neurological disease as multiparity was reported to improve outcomes after cerebral ischemia in female mice^81^. Transplant studies suggest novel therapeutic strategies, notably in cancer where a case report described regression of neuroblastoma after allogeneic hematopoietic cell transplantation^82^ and hematopoietic cell transplantation with microglia is being explored as treatment for advanced tumors^16,17^.

That microchimeric cells acquire many different fates suggests significant plasticity of integrating maternal cells likely with substantial input of the microenvironment on cell fate acquisition of the integrated cells. Our analytical methods allow better identification and characterization of microchimeric cells using high resolution transcriptomics data and open new lines of inquiry into understanding the biological phenomenon and consequences of Mc. We interpret these findings to indicate that the brain is not a privileged space with respect to Mc infiltration. Our data suggest that maternal cell migration may provide neurologic and immunologic advantage, as has been suggested by others^83^ and that integration of maternal microchimeric cells could be an important aspect of normal development. Together, the data show that naturally acquired Mc is prevalent and diverse in human brain, in a chronic brain disorder as well as in health, and that it persists in the human brain over the lifespan.

## Methods

### Ethics statement

The study included pediatric patients who received surgery as treatment for medication refractory epilepsy and whose parents/guardians consented for collection of research study specimens and data collection. The Internal Review Board (IRB) of the Fred Hutchinson Cancer Center (**FHCC**) approved this study (IRB file #7951, protocol 2648, RG1000938) and Seattle Children’s (IR file #11756), in accordance with institutional guidelines, the Declaration of Helsinki, and Title 45 United States Code of Federal Regulations, Part 46, Protection of Human Subjects.

### Selection of patients and acquisition of brain samples

Fifty-five medication refractory epilepsy subjects and their 55 mothers were recruited. The total number of study subjects was 178 as fathers (n=30) and siblings (n=38) were included when available. Buccal swabs were requested from participating family members. Brain specimens that were excess from clinically indicated surgeries were obtained from 53 patients. In one family sample was insufficient for genotyping so the final dataset available for microchimerism studies was 52 patient-mother pairs. In the final microchimerism dataset 23 patients were males and 29 females. Patients ages at surgery ranged from 28 days to 19 years, median 9.6 years. The most common epilepsy etiology was focal cortical dysplasia (**Supplemental Table 1**). Perinatal brain injury, hemorrhage, and vascular malformations were excluded. Brain specimens were flash frozen and kept at -80°C. Specimens were derived from hippocampus, frontal, parietal, occipital, and temporal lobes depending on the clinically indicated area for resection. Mothers were asked to complete a questionnaire for demographic and detailed reproductive history including any pregnancy complications and birth order of the patient.

#### DNA extraction

Genomic DNA was required for probands and their respective mothers to determine a genetic polymorphism unique to the mother to target for maternal Mc identification and quantification. DNA from maternal buccal swab samples was extracted using the QIAamp® DNA Mini Kit (QIAGEN, Valencia, CA) according to the manufacturer’s tissue protocol. To minimize contamination, DNA was extracted from brain tissues in a biosafety cabinet inside a clean room. The clean room suite was locked with limited access and consisted of a cascading set of positively pressurized rooms via HEPA filtered intakes with interlocked doors (such that only one door could be opened at once to maintain pressure). Extractions took place in the innermost room. The outermost room was a gowning room. The clean room was equipped with easily disinfected finishes and UV lights over benches, and all PCR reactions and storage of post-PCR was outside of the clean suite. ≤ 25 mg of frozen brain tissue was crushed until powder over liquid nitrogen with a previously autoclaved mortar and pestle and kept at -80°C, executed inside a cleaned biosafety cabinet (UV radiation, bleach 10%, ethanol 70%), at a rate of one sample per mortar/pestle. DNA was then extracted following the QIAamp® DNA Mini Kit (QIAGEN, Valencia, CA) according to the manufacturer’s tissue protocol, except the proteinase-K lysis step that was carried out overnight. DNA extractions were conducted in facilities physically separate from the ones in which Mc assays were conducted to avoid cross-contamination.

### Identification of genomic targets for Mc assay

Our laboratory developed an extensive panel of polymorphism-specific primers and fluorogenic probes to detect and quantify nonshared polymorphisms in highly sensitive real-time TaqMan quantitative polymerase chain reaction (qPCR). The qPCR assays include 22 targeting polymorphisms in HLA (historically chosen by our group because of marked polymorphism of HLA genes) and 18 for other non-HLA regions^19^. To identify a uniquely targetable maternal polymorphism not shared with the patient, HLA genotyping was first conducted for the patient and mother using a Luminex-based (One Lambda, Thermo Fisher Scientific Inc., Waltham, MA, U.S) polymerase chain reaction (PCR) sequence-specific oligonucleotide probe technique using beads and fluorescent labeling with analysis on a dual laser Luminex LX200 IS flow cytometer to genotype for *HLA-DRB1*, *-DRB3*, *-DRB4*, *-DRB5*, *-DQA1*, *-DQB1*, and *-B*. When a targetable HLA unique to the mother was unavailable, genotyping for non-HLA markers was performed using a multiplexed qPCR method previously established by our group^19^.

### Polymorphism-specific qPCR for Mc

After identifying a targetable polymorphism unique to the mother, Mc identification and quantification was performed with the appropriate TaqMan qPCR assay using the absolute quantification method. Each assay was designed to be highly sensitive (detecting Mc in the order of 1-in-a-million) and exclusively specific (never amplifying unintended targets)^19^. Each sample was measured in sextuplicates for the target assay and in duplicates for the reference assay in separate wells following previously published methods^21^. Total gDNA was calculated in genome equivalents (gEqs) via the reference standard curve and maternal gDNA via the target standard curve^21^. Maternal Mc was calculated as the ratio of these two quantities.

### Single-nucleus RNA barcoding and droplet-based capture

For a select number of probands with sufficiently leftover frozen brain tissue specimen, single nucleus whole RNA capture was performed using either the 10x Genomics Next GEM Single Cell 3’ v3.1 chemistry or the 10x Genomics GEM-X Single Cell 5’ v3 chemistry. For the generation of cell-barcoded complimentary DNA (cDNA) molecules, the appropriate 10x Genomics User Guide was followed (document #CG000317_Rev_B for the 3’ chemistry; #CG000733_Rev_A for the 5’ chemistry). Approximately 20–90 mg of frozen human brain tissue was used as input for the Minute™ Single Nucleus Isolation Kit for Neuronal Tissues/Cells (Invent Biotechnologies, Inc. Cat # BN-020). Nuclei were stained with trypan blue, counted, and concentration adjusted according to 10x protocol guidelines (approximately 1,000 nuclei/uL). About 10,000 nuclei were submitted for capture using the 10x Chromium Controller for Next GEM 3’ captures and approximately 20,000 nuclei were targeted for capture using the 10x Chromium X for GEM-X 5’ protocol. Following reverse transcription and cell barcoding in droplets, emulsions were broken and cDNA was purified using Dynal MyOne SILANE (ThermoFisher Scientific #37002D) followed by a PCR amplification (3’ v3.1 chemistry: 98°C for 3 min; 12 cycles of 98°C for 15 s, 63°C for 20 s, and 72°C for 1 min; 5’ v3 chemistry: 98°C for 45 s; 12 cycles of 98°C for 20 s, 63°C for 30 s, and 72°C for 1 min). Size selection was used to select the amplified cDNA molecules (> 800 bp) using SPRIselect reagent (Beckman Coulter #B23318). The majority of cDNA molecules in the final product had expected sizes of at least 600 bp and less than 2000 bp.

### Gene expression library construction and sequencing

For gene expression library construction, 100–250 ng (5’ capture chemistry) or between 25–150 ng (3’ capture chemistry) of amplified cDNA were enzymatically fragmented, end-repaired, A-tailed, and ligated to a partial TrueSeq adaptor. Additional Illumina adaptors and TrueSeq/TrueSeq sample indexes (P5, P7, i7 and i5) were added via PCR (98°C for 45 s; 14 cycles of 98°C for 20 s, 54°C for 30 s, and 72°C for 20 sec). Amplified DNA was size selected and purified with an expected size range between 300-600 bp. Libraries were sequenced on an Illumina NovaSeq 6000 sequencer using a pair-ended dual indexing configuration with 28 bp (Read1), 10 bp (i7 Index), 10 bp (i5 Index), 90 bp (Read2) for 3’ capture chemistry, and 26 bp (Read1), 10 bp (i7 Index), 10 bp (i5 Index), 90 bp (Read2) for 5’ capture chemistry, for a sequencing depth of ≥ 25,000 read pairs per cell.

### Single-nucleus gene expression data processing and quality control with Seurat

Illumina BCL files were demultiplexed and converted to FASTQ format using ‘bcl2fastq’ version 2.20.0 (Illumina, Inc.) through the cellranger-mkfastq function (10x Genomics, Inc.). Resulting FASTQ files were processed with ‘cellranger’ (versions 7.0.1 for the 3’-chemistry and 8.0.0 for the GEM-X 5’ chemistry) to align to the hg38 reference genome and count Gene Expression data. An output filtered feature barcode matrix file was then read on R (version 4.2.0) and converted into a ‘Seurat’ object^84^ (version 5.0.1). Initial quality control was used to subset out nuclei with too many (> 25,000) or too few (< 500) RNA counts and too many mitochondrial RNA counts (> 10% of a cell’s total RNA counts). Counts were then normalized (‘LogNormalize’ method, scale factor: 10,000), variable features were selected (‘vst’ selection method), features were centered and scaled (‘ScaleData()’ function), principal component analysis on the variable features was run, and UMAP (uniform manifold approximation and projection) low dimensional embedding^85^ and clustering using k-nearest-neighbor algorithm was performed on data. To remove multiplets, an artifact of single cell sequencing methods where two or more cells/nuclei receive the same barcode, ‘scrubletR’^28^ (version 0.2.1) was used on the RNA-seq count data in Seurat. scrubletR is an R implementation of the original Scrublet^28^ in Python. It simulates artificial doublets randomly from the captured data then assigns a doublet score (between 0 and 1) to each captured ‘cell’, based on the density of simulated doublets it co-localizes with, according to a k-nearest-neighbor classifier. Nuclei were considered doublets and removed if they had a doublet score > 0.2 after two consecutive scrubletR runs. To account for variations in single-cell/single-nucleus RNA-seq influenced by technical factors (i.e., samples run on different capture lanes, or experiments conducted in different times or laboratories), data integration was performed using ‘Harmony’ (version 1.2.3) to correct for batch effects^86^.

### Automatic cell type labeling using viewmastR

The cell types were automatically labeled using the ‘viewmastR’ package^32,87^ (version 0.2.1), a highly customizable classifier based on the Burn^88^ machine-learning/deep learning platform performing unsupervised classification of single cells, on the basis of known marker expression, between the query dataset and a well-annotated reference dataset using the ‘softmax’ regression method, relying on the top 10,000 genes commonly variable across the datasets. In this study using single-nucleus from human brain biopsies, the reference dataset was the well annotated Allen Brain Map Human M1 Cortex single-nucleus capture Atlas of post-mortem human brain specimens surveying cell type diversity in the primary motor cortex^33,89^.

### Genetic demultiplexing of allogeneic entities with *cellector* and *souporcell*

To perform variant calling and distinguish allogeneic cells (arising from naturally acquired Microchimerism) vs. autochthonous cells in the human brain biopsies, both ‘cellector’^20^ (version 1.0.0) and ‘souporcell’^29^ (version ≥ 2.0; Singularity version 3.5.3) were used. *Souporcell* is a genetic demultiplexing method that clusters cells by genotype using genetic variants detected in scRNA-seq reads^29^, whereas *cellector* was designed specifically to identify rare cells with a distinct genotype in highly imbalanced mixtures of genotype populations. *Cellector* models the majority genotype using beta-binomial distributions and computes a cell-level log-likelihood normalized by the number of informative loci. Cells with normalized log-likelihoods below a lower outlier threshold of Q_1_ –m × IQR are provisionally excluded from the majority population (default: m = 5), the model is recomputed, and this procedure is iterated until the set of excluded cells converges^90^. Both methods were supplied with cellranger-generated BAM files aligned to GRCh38, filtered cell-barcode files, the corresponding GRCh38 reference FASTA, and a VCF containing common variants from the 1000 Genomes Project. For *souporcell*, ‘--skip_remap’ and ‘--ignorè were enabled, and the expected number of genotype clusters was specified using ‘-k’. The sparse mixture-model output from *souporcell* was log-normalized and visualized according to genotype assignment.

### HLA haplotype phasing from host- and microchimeric-cell BAMs

Clusters of variant haplotypes within HLA genes were reconstructed independently from BAM files containing reads assigned to host or microchimeric cells using a custom ‘R’ workflow (version 4.2.0). GENCODE v36 gene annotations were imported for the extended MHC region on chromosome 6. For each HLA or HLA-associated gene, a representative transcript was selected by prioritizing MANE Select, Ensembl canonical, protein-coding, APPRIS principal, and basic transcripts, followed by total annotated exon length. Primary mapped alignments overlapping the full genomic span of the selected transcript were extracted with ‘Rsamtools’ (version 2.14.0); unmapped, secondary, and supplementary alignments were excluded. Read sequences were projected onto reference coordinates from their CIGAR strings, retaining substitutions, deletions, and insertions.

Each transcript span was surveyed using 600-bp windows with 300-bp overlap. Windows containing at least five host reads and one Mc read were ranked primarily by Mc read depth and then by host read depth. Within each window, positions covered by at least five host reads or two Mc reads were retained, without applying a minor-allele-frequency threshold. Reads covering at least five retained positions were clustered by a categorical k-modes procedure. Briefly, candidate read haplotypes were initialized from randomly selected reads, reads were assigned to the haplotype with the lowest mismatch rate across shared positions, and haplotypes were updated using the modal base at each position. Models containing two to six candidate haplotypes were evaluated using five random initializations, and the solution minimizing the total number of read–haplotype mismatches was retained.

Haplotypes representing at least 10% of assigned reads were considered supported, with a maximum of two retained haplotypes per host or Mc population. Supported Mc and host haplotypes were compared across positions observed in both sequences, and the number and proportion of mismatching bases were recorded.

### Mc vs. host differential expression analysis

One approach was label-agnostic (i.e., independent of any inferred cell identity). Each microchimeric cell/nucleus was matched to its 30 nearest host cells/nuclei from the same brain using Euclidean distance on the first 30 principal components. For each gene, normalized RNA expression in the Mc cell was compared with the mean expression among its matched host neighbors to obtain a local Mc–host expression difference (Δ expression). Differences were first averaged within each biological brain to avoid disproportionate weighting of individuals containing larger numbers of microchimeric cells. For epilepsy samples generated using both 3′ and 5′ capture chemistries, or within the same capture chemistry but done on different brain regions, capture-specific estimates from the same individual were combined at the proband level with equal weighting. Healthy and epilepsy cohorts were analyzed independently and gene-level effects were summarized using the median proband-level Δ expression. Directional consistency was additionally assessed across probands, requiring concordant expression changes in ≥ 80% of healthy brains or epilepsy brains for cohort-specific classification.

Another approach was cell-label-supervised, conducted within predefined brain cell types. Cells/nuclei were first stratified according to their viewmastR-predicted identity and Mc status. Cell types represented by microchimeric cells in at least two independent probands were analyzed, providing biological replication for the paired differential-expression model. Within each Mc-positive proband and capture, all Mc cells of a given cell type were retained and an equal number of host cells of the same viewmastR class were randomly sampled, thereby permitting inclusion of probands containing only a single microchimeric cell while avoiding the large imbalance in cell numbers between Mc and host populations. Raw RNA counts were summed to generate paired Mc and host pseudobulks for each biological proband; where a proband contributed multiple informative captures, capture-level counts were combined after balancing. Differential expression was tested using an edgeR quasi-likelihood model^35,36^ including proband as a blocking factor: ‘∼ proband + Mc status’ (‘edgeR’ version 3.40.2). Host-cell sampling was repeated 300 times, and gene-level effects were summarized by the median log-fold change across iterations, and the proportion of iterations showing the same direction of effect.

### Hallmark gene set enrichment analysis

Gene set enrichment analysis (GSEA) was performed separately for healthy and epilepsy cohorts using genes ranked by their mean proband-level matched expression difference between microchimeric cells and their 30 nearest host-cell neighbors. Hallmark gene sets from the Molecular Signatures Database^91^ (‘msigdbr’; version 24.1.0) were analyzed using preranked GSEA with ‘fgsea::fgseaMultilevel’^92^ (version 1.22.0), restricting analysis to gene sets containing 15–500 represented genes. Enrichment was summarized by the normalized enrichment score (NES) with positive values indicating pathways enriched among genes expressed more highly in microchimeric cells and negative values indicating enrichment among genes expressed less highly in microchimeric cells relative to matched host cells. P values were adjusted for multiple testing using the Benjamini-Hochberg false discovery rate (FDR), with FDR <0.05 considered significant and FDR <0.10 considered suggestive in this exploratory analysis.

### Analysis of publicly available datasets for microchimerism

We applied our analytical pipeline to publicly available snRNAseq datasets from Herring et al. (GSE168408)^21^, cognitively normal controls from Lau et al. (GSE157827),^22^ and healthy controls from Feng et al. (GSE301908).^23^ All datasets were generated with 10x Genomics technology, included raw FASTQ files, and contained appropriate sample metadata such as age and sex. Data were retrieved from the Sequence Read Archive (**SRA**) using the SRA Toolkit (v3.0.0) and then processed with ‘cellranger’ to align to the hg38 reference genome and count Gene Expression data as described above in the section ‘*Single-nucleus gene expression data processing and quality control with Seurat’.* Integration was performed using Harmony to correct for batch effects. We also used viewmastR to annotate cell types and *cellector* to identify microchimeric cells, both as described in relevant sections above.

## ACKNOWLEDGEMENTS

This research was supported by the National Institutes of Health grants KL2 TR002317 (AM), R21 NS071418 (JLN) and R21 NS118249 (JLN and JRW). We additionally thank Dr. David Schechner at UW for use of his Hyak slice for assembling the public datasets. The authors are grateful to the participants who consented to participate in the study.

## CONTRIBUTIONS

JLN conceived the study, SBK, AM, JRW and JLN designed the experiments. JR conceived a key aspect of the study design and contributed comments for the paper. JO contributed to the study design, analyses, and comments for the paper. SBK, AM, CG and FU performed the experiments. DTAE, DG, JRW, and JLN provided platforms and resources to conduct experiments. SBK, AM, HH, RB, and SNF contributed key elements into the computational analysis. SBK, AM, JRW and JLN wrote the paper. All authors have reviewed the manuscript.

## COMPETING INTERESTS

All authors declare no competing interests.

## FIGURE AND TABLE LEGENDS

**Supplementary Figure S1. Expressed variant loci contributing to cellector minority-genotype assignments.** Per-locus summaries are shown for all 56 snRNA-seq captures: 15 epilepsy captures from the BRE cohort and 41 control captures. Each facet represents one capture and shows up to 20 displayed loci from the cellector locus-contribution output. For each locus, the alternate-allele frequency in the cellector minority/anomaly group is plotted to the left of zero and that in the majority/host group to the right; bar length gives the absolute allele frequency. Bar color encodes the log likelihood of the minority group normalized per cell. Loci are labeled by GRCh38 genomic coordinate. BRE, Brain Epilepsy; snRNA-seq, single-nucleus RNA sequencing.

**Supplementary Figure S2. HLA-transcript detection relative to the global transcriptome.** Each point represents one gene, plotted according to the percentage of cells or nuclei with detectable expression and its aggregate counts per million (CPM), shown on a logarithmic scale. HLA genes are highlighted and labeled in red. From top to bottom, plots show 11 BRE 3′ brain-nucleus captures, four BRE 5′ brain-nucleus captures, 41 downloaded control brain-nucleus datasets, and a downloaded 5′ single-cell bone-marrow mononuclear-cell (BMMC) dataset (sample “AML_402” from https://doi.org/10.1101/2025.09.12.675951 [Ref. 31]). Compared with the capture as whole single cells, HLA transcripts were detected in a diminished proportion of nuclei in the brain datasets, limiting the read coverage available for HLA haplotype phasing. BMMC, bone-marrow mononuclear cell; BRE, Brain Epilepsy; CPM, counts per million; HLA, human leukocyte antigen.

**Supplementary Figure S3. Read-backed HLA haplotype phasing of host and microchimeric nuclei.** Haplotype clusters were reconstructed independently from reads assigned to host and candidate microchimeric nuclei within selected HLA-gene windows. Rows show supported host or Mc haplotypes; parentheses give the number and percentage of assigned reads represented by each haplotype cluster. **A,** Output in which a Mc haplotype matched one host haplotype across mutually observed positions. **B,** Output in which a Mc haplotype was discordant with the host haplotype or haplotypes covered within the displayed window. **C,** Uninformative output in which limited variant availability or overlap between host and Mc reads precluded classification as shared or nonshared. Gray tiles denote matching positions, discordant bases are colored by nucleotide (A, magenta; C, cyan; G, green; T, orange), and white denotes missing coverage. Coordinates refer to GRCh38-3.0.0. HLA, human leukocyte antigen; Mc, microchimerism. **For “Herring-14yo HLA−B”, Mc_hap1 and Mc_hap2 are identical except at a single position, backed by a single read in haplotype Mc_hap2; if sequencing error is to be considered, Mc_hap2 may be dismissed and only Mc_hap1 considered.

**Supplementary Figure S4. Cell-type composition of epilepsy and healthy snRNA-seq captures.** Stacked bars show the relative abundance of viewmastR-assigned cell types among all quality-controlled nuclei in each capture; each bar sums to 1. Cell-type assignments used the Allen Brain Map Human M1 10x atlas as reference. **A,** Eleven BRE 3′ captures and four BRE 5′ captures, representing 10 epilepsy probands. Repeated proband labels indicate different brain regions or capture chemistries. **B,** Forty-one healthy captures comprising 29 Herring, 9 Lau, and 3 Feng probands, ordered broadly by age. Annotation strips indicate proband sex and sampled brain region. Astro, astrocyte; Chodl, chondrolectin-expressing; CT, corticothalamic; Endo, endothelial; ET, extratelencephalic; IT, intratelencephalic; L, cortical layer; Lamp5, LAMP5-expressing; Micro-PVM, microglia/perivascular macrophage; NP, near-projecting; Oligo, oligodendrocyte; OPC, oligodendrocyte precursor cell; Pvalb, Parvalbumin-positive; Sst, Somatostatin-expressing; Vip, vasoactive intestinal polypeptide-expressing; VLMC, vascular and leptomeningeal cell.

**Supplementary Figure S5. Hallmark gene-set enrichment in microchimeric nuclei relative to matched host nuclei.** Genes were ranked using Mc-host normalized-expression differences obtained by matching each candidate microchimeric nucleus to its 30 nearest host nuclei from the same proband. Analyses were performed separately for 6 Mc-positive epilepsy probands and 28 Mc-positive control probands. Each point represents one Hallmark gene set; the x axis shows the normalized enrichment score (NES) in healthy and the y axis the NES in epilepsy. Positive NES values indicate enrichment among genes expressed more highly in microchimeric nuclei, whereas negative values indicate enrichment among genes expressed less highly in microchimeric nuclei relative to matched host nuclei. Dashed lines mark NES = 0, and the dotted diagonal marks equal NES values in the two cohorts. Green denotes gene sets flagged at the exploratory FDR < 0.10 threshold in the figure, whereas gray denotes other gene sets; no pathway met the conventional FDR < 0.05 threshold. FDR, false discovery rate; Mc, microchimeric; NES, normalized enrichment score.

**Supplementary Figure S6. Capture-level abundance and cell-type composition of microchimeric nuclei in epilepsy brain and comparison with maternal-specific qPCR. A,** Mc frequency across 15 epilepsy captures from 10 probands, calculated as cellector-identified microchimeric nuclei divided by all nuclei and shown on a log10 scale. Mc was detected in eight captures. Pie-chart segments show the viewmastR-assigned cell-type composition of candidate microchimeric nuclei; black circles indicate captures in which microchimeric nuclei were not detected. **B,** Capture-level comparison of Mc proportions measured by maternal-polymorphism-specific qPCR and source-agnostic cellector classification. Horizontal red and vertical purple error bars show 95% Wilson-score confidence intervals for qPCR and cellector estimates, respectively; the blue dashed line is the identity line. Mc, microchimerism; qPCR, quantitative polymerase chain reaction.

**Supplementary Figure S7. Total UMI-count distributions in microchimeric and host nuclei.** Violin plots compare pooled distributions of log10-transformed total UMI counts per nucleus between cellector-classified microchimeric nuclei (red) and host nuclei (gray) in the BRE 3′, BRE 5′, and control datasets. Violin width represents nucleus density. BRE, Brain Epilepsy; Mc, microchimerism; UMI, unique molecular identifier.

**Supplementary Table S1. Clinical and specimen characteristics of BRE probands.** Brain regions, sex, age at seizure onset, age at resection, and pathologic diagnosis are reported for probands with surgically resected epilepsy tissue. Gray shading identifies probands investigated by snRNA-seq. BRE, Brain Epilepsy; snRNA-seq, single-nucleus RNA sequencing.

**Supplementary Table S2. Sequencing and quality-control metrics for BRE single-nucleus RNA-sequencing captures.** Rows represent 15 capture libraries from 10 epilepsy probands and report brain region, pre-quality-control nuclei and sequencing metrics, gene and UMI counts, filtering thresholds, final retained nuclei, percentage retained, and 10x Genomics capture chemistry. Repeated proband IDs indicate distinct brain regions or capture chemistries. Gray shading identifies GEM-X 5′ libraries; unshaded rows identify Next GEM 3′ libraries. BRE, Brain Epilepsy; GEO, Gene Expression Omnibus; QC, quality control; snRNA-seq, single-nucleus RNA sequencing; UMI, unique molecular identifier.

## Notes

### Competing Interest Statement

The authors have declared no competing interest.

### Summary of Updates

We have added a substantial amount of new data and analyses to the manuscript draft and reorganized the presentation of data. Figure 1 has been split into two figures with expanded analyses added to both. Figure 2 has become Figure 3 and we have added additional panels with more analyses. Likewise we have expanded our Supplemental Figures. The manuscript text has been updated to reflect these additions.

## REFERENCES

1. Nelson, J. L. & Lambert, N. C. The when, what, and where of naturally-acquired microchimerism. Semin. Immunopathol. 47, 20 (2025).

2. Kinder, J. M., Stelzer, I. A., Arck, P. C. & Way, S. S. Immunological implications of pregnancy-induced microchimerism. Nat. Rev. Immunol. 17, 483–494 (2017).

3. Loubiere, L. S. et al. Maternal microchimerism in healthy adults in lymphocytes, monocyte//macrophages and NK cells. Lab Invest 86, 1185–1192 (2006).

4. Stevens, A., Hermes, H., Rutledge, J., Buyon, J. & Nelson, J. Myocardial-tissue-specific phenotype of maternal microchimerism in neonatal lupus congenital heart block. Lancet 362, 1617–1623 (2003).

5. Srivatsa, B. et al. Microchimerism of presumed fetal origin in thyroid specimens from women: a case-control study. Lancet 358, 2034–2038 (2001).

6. Nelson, J. et al. Maternal microchimerism in peripheral blood in type 1 diabetes and pancreatic islet beta cell microchimerism. Proc Natl Acad Sci 104, 1637–1642 (2007).

7. Stevens, A. M., Hermes, H. M., Kiefer, M. M., Rutledge, J. C. & Nelson, J. L. Chimeric Maternal Cells with Tissue-Specific Antigen Expression and Morphology are Common in Infant Tissues. Pediatr. Dev. Pathol. Off. J. Soc. Pediatr. Pathol. Paediatr. Pathol. Soc. 12, 337–346 (2009).

8. Khosrotehrani, K. et al. Presence of chimeric maternally derived keratinocytes in cutaneous inflammatory diseases of children: the example of pityriasis lichenoides. J. Invest. Dermatol. 126, 345–348 (2006).

9. Jonsson, A. M., Uzunel, M., Götherström, C., Papadogiannakis, N. & Westgren, M. Maternal microchimerism in human fetal tissues. Am. J. Obstet. Gynecol. 198, 325.e1–325.e6 (2008).

10. Snethen, H., Ye, J., Gillespie, K. M. & Scolding, N. J. Maternal micro-chimeric cells in the multiple sclerosis brain. Mult. Scler. Relat. Disord. 40, 101925 (2020).

11. Chan, W. F. N. et al. Male Microchimerism in the Human Female Brain. PLoS ONE 7, (2012).

12. Broestl, L., Rubin, J. B. & Dahiya, S. Fetal microchimerism in human brain tumors. Brain Pathol. Zurich Switz. 28, 484–494 (2018).

13. Zeng, X. X. et al. Pregnancy-associated progenitor cells differentiate and mature into neurons in the maternal brain. Stem Cells Dev. 19, 1819–1830 (2010).

14. Fujimoto, K. et al. Whole-embryonic identification of maternal microchimeric cell types in mouse using single-cell RNA sequencing. Sci. Rep. 12, 18313 (2022).

15. Schepanski, S. et al. Pregnancy-induced maternal microchimerism shapes neurodevelopment and behavior in mice. Nat Commun 13, 4571 (2022).

16. Shibuya, Y. et al. Treatment of a genetic brain disease by CNS-wide microglia replacement. Sci. Transl. Med. 14, eabl9945 (2022).

17. Loeb, A. M., Pattwell, S. S., Meshinchi, S., Bedalov, A. & Loeb, K. R. Donor bone marrow-derived macrophage engraftment into the central nervous system of patients following allogeneic transplantation. Blood Adv 7, 5851–5859 (2023).

18. Lambert, N. C. et al. Quantification of maternal microchimerism by HLA-specific real-time polymerase chain reaction: studies of healthy women and women with scleroderma. Arthritis Rheum 50, 906–14 (2004).

19. Kanaan, S. B., Urselli, F., Radich, J. P. & Nelson, J. L. Ultrasensitive chimerism enhances measurable residual disease testing after allogeneic hematopoietic cell transplantation. Blood Adv 7, 6066–6079 (2023).

20. Heaton, H. et al. Cellector: A tool to detect foreign genotype cells in scRNAseq data with applications in leukemia and microchimerism. 2026.03.26.714571 Preprint at 10.64898/2026.03.26.714571 (2026).

21. Herring, C. A. et al. Human prefrontal cortex gene regulatory dynamics from gestation to adulthood at single-cell resolution. Cell 185, 4428–4447 e28 (2022).

22. Lau, S.-F., Cao, H., Fu, A. K. Y. & Ip, N. Y. Single-nucleus transcriptome analysis reveals dysregulation of angiogenic endothelial cells and neuroprotective glia in Alzheimer’s disease. Proc. Natl. Acad. Sci. U. S. A. 117, 25800–25809 (2020).

23. Feng, R. et al. Single-cell spatial transcriptomic profiling defines a pathogenic inflammatory niche in chronic active multiple sclerosis lesions. Immunity 58, 2989–3005.e10 (2025).

24. Shao, T.-Y. et al. Reproductive outcomes after pregnancy-induced displacement of preexisting microchimeric cells. Science 10.1126/science.adf9325 (2023) doi:10.1126/science.adf9325.

25. Boddy, A. M., Fortunato, A., Sayres, M. W. & Aktipis, A. Fetal microchimerism and maternal health: A review and evolutionary analysis of cooperation and conflict beyond the womb. BioEssays 37, 1106–1118 (2015).

26. Haig, D. Does microchimerism mediate kin conflicts? Chimerism 5, (2014).

27. Ortolano, N. The neXt Generation of Single Cell RNA-Seq: An Introduction to GEM-X Technology. 10x Genomics (2024).

28. Wolock, S. L., Lopez, R. & Klein, A. M. Scrublet: Computational Identification of Cell Doublets in Single-Cell Transcriptomic Data. Cell Syst 8, 281–291 e9 (2019).

29. Heaton, H. et al. Souporcell: robust clustering of single-cell RNA-seq data by genotype without reference genotypes. Nat Methods 17, 615–620 (2020).

30. Horton, R. et al. Gene map of the extended human MHC. Nat. Rev. Genet. 5, 889–899 (2004).

31. Kanaan, S. B. et al. Quantifying HLA transcripts by genotype in chimeric mixtures at single-cell resolution. 2025.09.12.675951 Preprint at 10.1101/2025.09.12.675951 (2025).

32. Krakow, E. F. et al. HA-1–targeted T-cell receptor T-cell therapy for recurrent leukemia after hematopoietic stem cell transplantation. Blood 144, 1069–1082 (2024).

33. Human M1 10x. https://brain-map.org/our-research/cell-types-taxonomies/cell-types-database-rna-seq-data/human-m1-10x.

34. van Tilborg, E. et al. Origin and dynamics of oligodendrocytes in the developing brain: Implications for perinatal white matter injury. Glia 66, 221–238 (2018).

35. Robinson, M. D., McCarthy, D. J. & Smyth, G. K. edgeR: a Bioconductor package for differential expression analysis of digital gene expression data. Bioinformatics 26, 139–140 (2010).

36. Chen, Y., Lun, A. T. L. & Smyth, G. K. From reads to genes to pathways: differential expression analysis of RNA-Seq experiments using Rsubread and the edgeR quasi-likelihood pipeline. Preprint at 10.12688/f1000research.8987.2 (2016).

37. Gehman, L. T. et al. The splicing regulator Rbfox1 (A2BP1) controls neuronal excitation in the mammalian brain. Nat. Genet. 43, 706–711 (2011).

38. Kim, S., Shin, J. J., Kang, M., Yi, Y. & Kim, E. Synaptic and Non-Synaptic Functions of PTPRD: A Receptor Tyrosine Phosphatase at the Crossroads of Neural Circuitry and Metabolism. J. Neurochem. 169, e70292 (2025).

39. Bogaciu, C. A. & Rizzoli, S. O. Membrane trafficking of synaptic adhesion molecules. J. Physiol. 603, 5921–5934 (2025).

40. Strehlow, V., Myers, K. A., Morgan, A. T., Scheffer, I. E. & Lemke, J. R. GRIN2A-Related Disorders. in GeneReviews® (eds Adam, M. P., Bick, S., Mirzaa, G. M., Wallace, S. E. & Amemiya, A.) (University of Washington, Seattle, Seattle (WA), 1993).

41. Gordon-Weeks, P. R. & Fournier, A. E. Neuronal cytoskeleton in synaptic plasticity and regeneration. J. Neurochem. 129, 206–212 (2014).

42. Widagdo, J. & Anggono, V. The m6A-epitranscriptomic signature in neurobiology: from neurodevelopment to brain plasticity. J. Neurochem. 147, 137–152 (2018).

43. Barry, G. et al. The long non-coding RNA NEAT1 is responsive to neuronal activity and is associated with hyperexcitability states. Sci. Rep. 7, 40127 (2017).

44. Bezerra, G. A. et al. Structure of human dipeptidyl peptidase 10 (DPPY): a modulator of neuronal Kv4 channels. Sci. Rep. 5, 8769 (2015).

45. Henley, J. M. et al. Kainate and AMPA receptors in epilepsy: Cell biology, signalling pathways and possible crosstalk. Neuropharmacology 195, 108569 (2021).

46. González-Velasco, O., Papy-García, D., Le Douaron, G., Sánchez-Santos, J. M. & De Las Rivas, J. Transcriptomic landscape, gene signatures and regulatory profile of aging in the human brain. Biochim. Biophys. Acta BBA - Gene Regul. Mech. 1863, 194491 (2020).

47. Wu, J. et al. Targeting MEF2A suppresses microglial hyperactivation and synaptic phagocytosis to attenuate epilepsy pathogenesis. Cell Death Dis. 17, 645 (2026).

48. Ip, J. P. K., Fu, A. K. Y. & Ip, N. Y. CRMP2: Functional Roles in Neural Development and Therapeutic Potential in Neurological Diseases. The Neuroscientist 20, 589–598 (2014).

49. Südhof, T. C. Towards an Understanding of Synapse Formation. Neuron 100, 276–293 (2018).

50. Mezey, É. et al. Transplanted bone marrow generates new neurons in human brains. Proc. Natl. Acad. Sci. 100, 1364–1369 (2003).

51. Guthrie, K. A. et al. Statistical Methods for Unusual Count Data: Examples From Studies of Microchimerism. Am. J. Epidemiol. 184, 779–786 (2016).

52. Lo, E. S., Lo, Y. M., Hjelm, N. M. & Thilaganathan, B. Transfer of nucleated maternal cells into fetal circulation during the second trimester of pregnancy. Br. J. Haematol. 100, 605–606 (1998).

53. Aydın, M. Ş., Yiğit, E. N., Vatandaşlar, E., Erdoğan, E. & Öztürk, G. Transfer and Integration of Breast Milk Stem Cells to the Brain of Suckling Pups. Sci. Rep. 8, 14289 (2018).

54. Borges, A., Castellan, F. & Irie, N. Emergent roles of maternal microchimerism in postnatal development. Dev. Growth Differ. 65, 75–81 (2023).

55. Harry, G. J. Microglia during development and aging. Pharmacol. Ther. 139, 313–326 (2013).

56. Harris, J., Tomassy, G. S. & Arlotta, P. Building blocks of the cerebral cortex: from development to the dish. WIREs Dev. Biol. 4, 529–544 (2015).

57. Yao, Z. et al. A taxonomy of transcriptomic cell types across the isocortex and hippocampal formation. Cell 184, 3222–3241.e26 (2021).

58. Wang, L. et al. Molecular and cellular dynamics of the developing human neocortex. Nature 647, 169–178 (2025).

59. Fisher, J. et al. Cortical somatostatin long-range projection neurons and interneurons exhibit divergent developmental trajectories. Neuron 112, 558–573.e8 (2024).

60. DeSisto, J. et al. Single-Cell Transcriptomic Analyses of the Developing Meninges Reveal Meningeal Fibroblast Diversity and Function. Dev. Cell 54, 43–59.e4 (2020).

61. Martínez-Cerdeño, V. et al. Behavior of Xeno-Transplanted Undifferentiated Human Induced Pluripotent Stem Cells Is Impacted by Microenvironment Without Evidence of Tumors. Stem Cells Dev. 26, 1409–1423 (2017).

62. Ricardo C. H. del Rosario et al. Sibling chimerism among microglia in marmosets. bioRxiv 2023.10.16.562516 (2023) doi:10.1101/2023.10.16.562516.

63. Villeda, S. A. et al. The ageing systemic milieu negatively regulates neurogenesis and cognitive function. Nature 477, 90–94 (2011).

64. Villeda, S. A. et al. Young blood reverses age-related impairments in cognitive function and synaptic plasticity in mice. Nat. Med. 20, 659–663 (2014).

65. Tan, X.-W. et al. Fetal Microchimerism in the Maternal Mouse Brain: A Novel Population of Fetal Progenitor or Stem Cells Able to Cross the Blood-Brain Barrier? Stem Cells 23, 1443–1452 (2005).

66. Sbeih, M. et al. CCL2 recruits fetal microchimeric cells and dampens maternal brain damage in post-partum mice. Neurobiol. Dis. 174, 105892 (2022).

67. Castellan, F. S. & Irie, N. Postnatal depletion of maternal cells biases T lymphocytes and natural killer cells’ profiles toward early activation in the spleen. Biol. Open 11, (2022).

68. Stelzer, I. A. et al. Vertically transferred maternal immune cells promote neonatal immunity against early life infections. Nat. Commun. 12, 1–14 (2021).

69. Harrington, W. E. et al. Maternal Microchimerism Predicts Increased Infection but Decreased Disease due to Plasmodium falciparum During Early Childhood. J. Infect. Dis. 215, 1445–1451 (2017).

70. Kinder, J. M. et al. Cross-Generational Reproductive Fitness Enforced by Microchimeric Maternal Cells. Cell 162, 505–515 (2015).

71. Leveque, L. & Khosrotehrani, K. Can maternal microchimeric cells influence the fetal response toward self antigens? Chimerism 2, 71–77 (2011).

72. Wrenshall, L. E., Stevens, E. T., Smith, D. R. & Miller, J. D. Maternal microchimerism leads to the presence of interleukin-2 in interleukin-2 knock out mice: Implications for the role of interleukin-2 in thymic function. Cell. Immunol. 245, 80–90 (2007).

73. Berry, S. M. et al. Association of maternal histocompatibility at Class II loci with maternal microchimerism in the fetus. Pediatr Res 56, 73–78 (2004).

74. Guettier, C. et al. Male cell microchimerism in normal and diseased female livers from fetal life to adulthood. Hepatology 42, 35–43 (2005).

75. Kumar, P. et al. Single-cell transcriptomics and surface epitope detection in human brain epileptic lesions identifies pro-inflammatory signaling. Nat. Neurosci. 25, 956–966 (2022).

76. Kaplan, J. & Land, S. Influence of Maternal-Fetal Histocompatibility and MHC Zygosity on Maternal Microchimerism. J Immunol 174, 7123–7128 (2005).

77. Nordahl, C. W. et al. Maternal autoantibodies are associated with abnormal brain enlargement in a subgroup of children with autism spectrum disorder. Brain. Behav. Immun. 30, 61–65 (2013).

78. Fox-Edmiston, E. & Van de Water, J. Maternal Anti-Fetal Brain IgG Autoantibodies and Autism Spectrum Disorder: Current Knowledge and its Implications for Potential Therapeutics. CNS Drugs 29, 715–724 (2015).

79. Hemon, M. et al. Microchimeric cells promote production of rheumatoid arthritis-specific autoantibodies. J. Autoimmun. 146, 103238 (2024).

80. Prüss, H. Autoantibodies in neurological disease. Nat. Rev. Immunol. 21, 798–813 (2021).

81. Ritzel, R. M., et al. Multiparity improves outcomes after cerebral ischemia in female mice despite features of increased metabovascular risk. Proc Natl Acad Sci U A 114, E5673–E5682 (2017).

82. Inoue, M. et al. Graft-versus-tumor effect in a patient with advanced neuroblastoma who received HLA haplo-identical bone marrow transplantation. Bone Marrow Transplant. 32, 103–106 (2003).

83. Slaats, E. et al. Maternal microchimeric cell trafficking and its biological consequences depend on the onset of inflammation at the feto-maternal interface. Semin. Immunopathol. 47, 8 (2025).

84. Hao, Y. et al. Integrated analysis of multimodal single-cell data. Cell 184, 3573–3587 e29 (2021).

85. Becht, E. et al. Dimensionality reduction for visualizing single-cell data using UMAP. Nat Biotechnol 10.1038/nbt.4314 (2018) doi:10.1038/nbt.4314.

86. Korsunsky, I. et al. Fast, sensitive and accurate integration of single-cell data with Harmony. Nat. Methods 16, 1289–1296 (2019).

87. Furlan, S. viewmastR: Automated single-cell genomic cell type assignment. R package version 0.2.3 (2024).

88. Simard, N., Fortier-Dubois, L., Tadjibaev, D., Lagrange, G., & Burn Framework Contributors. Burn. (2026).

89. Bakken, T. E. et al. Comparative cellular analysis of motor cortex in human, marmoset and mouse. Nature 598, 111–119 (2021).

90. Heaton, H. wheaton5/cellector: statistical model for finding anomalous genotype cells in mixed genotype scRNAseq data. GitHub https://github.com/wheaton5/cellector.

91. Liberzon, A. et al. Molecular signatures database (MSigDB) 3.0. Bioinformatics 27, 1739–1740 (2011).

92. Korotkevich, G. et al. Fast gene set enrichment analysis. 060012 Preprint at 10.1101/060012 (2021).

93. Edgar, R., Domrachev, M. & Lash, A. E. Gene Expression Omnibus: NCBI gene expression and hybridization array data repository. Nucleic Acids Res. 30, 207–210 (2002).

