## Supplemental Figures 1-7 for "Microchimerism in the Human Brain, Quantitative Assessment and Single Nucleus Profiling Establish Cell Types and Diversity"

### SUPPLEMENTAL FIGURE 1

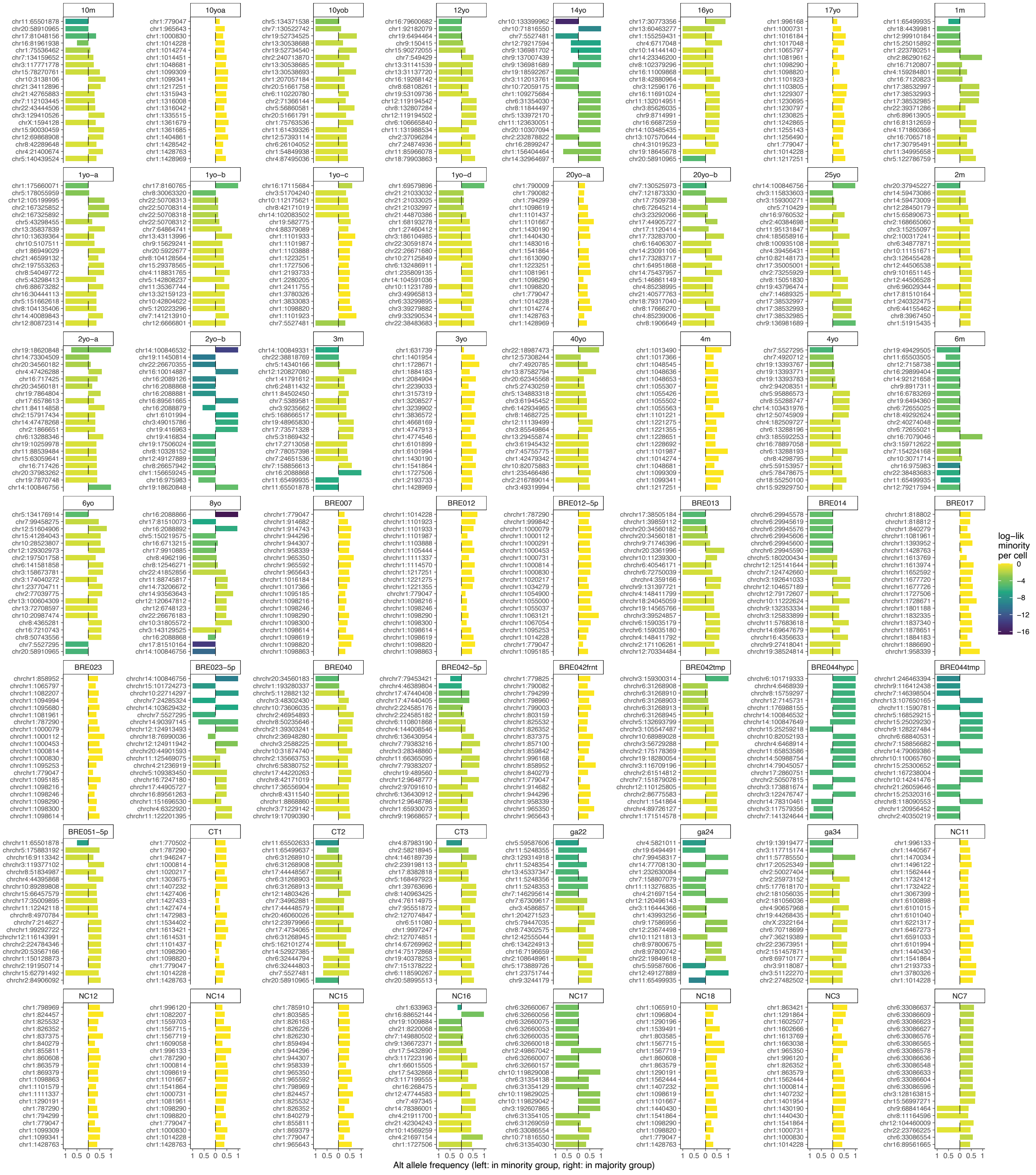

**SUPPLEMENTAL FIGURE 2**

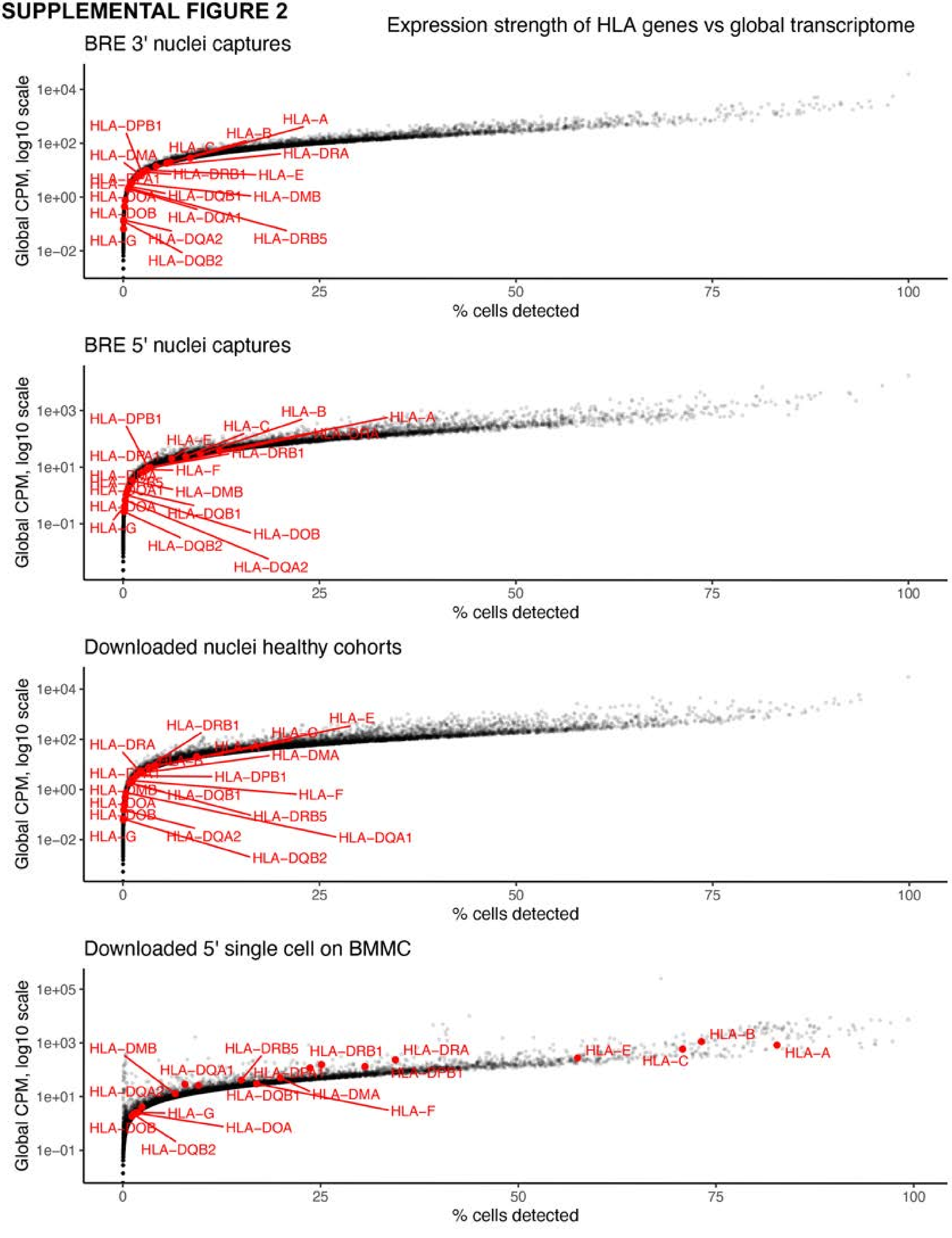

### SUPPLEMENTAL FIGURE 3

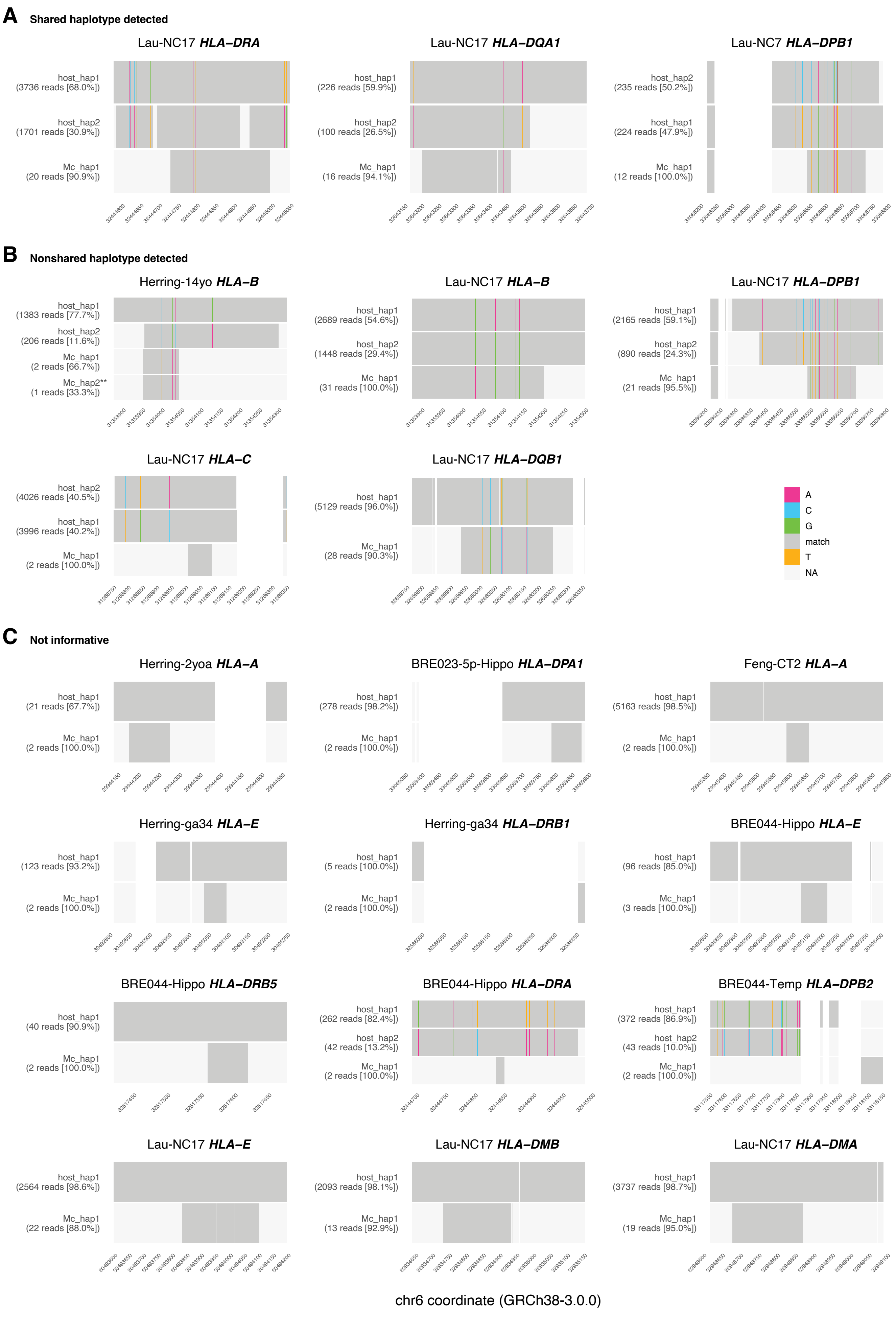

### SUPPLEMENTAL FIGURE 4

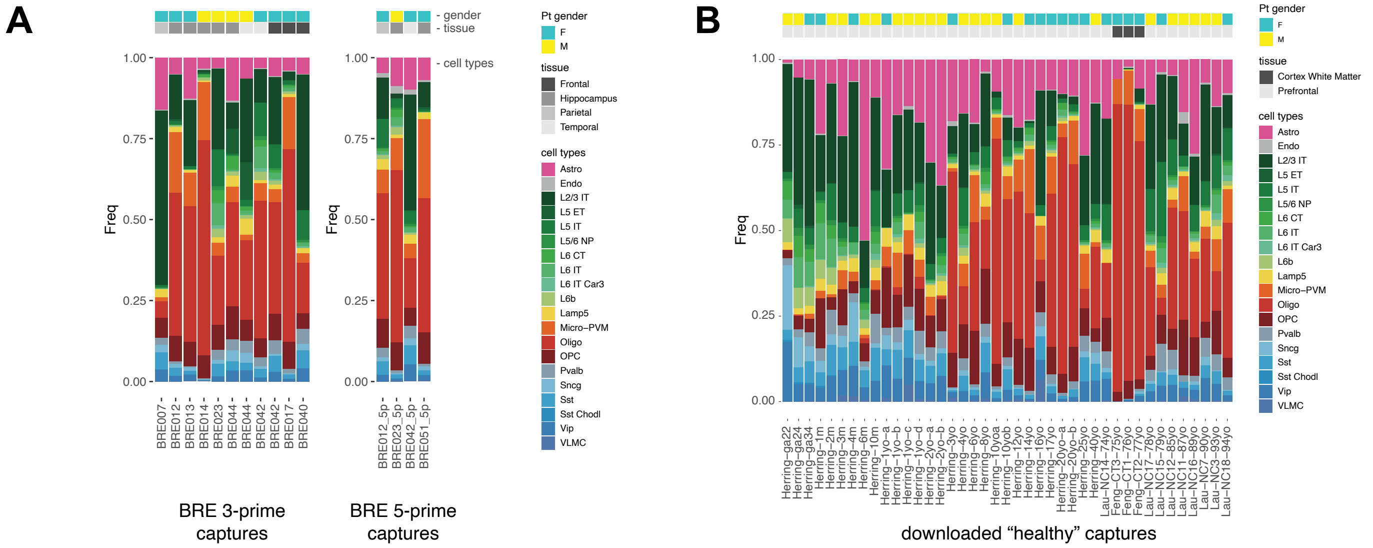

### SUPPLEMENTAL FIGURE 5

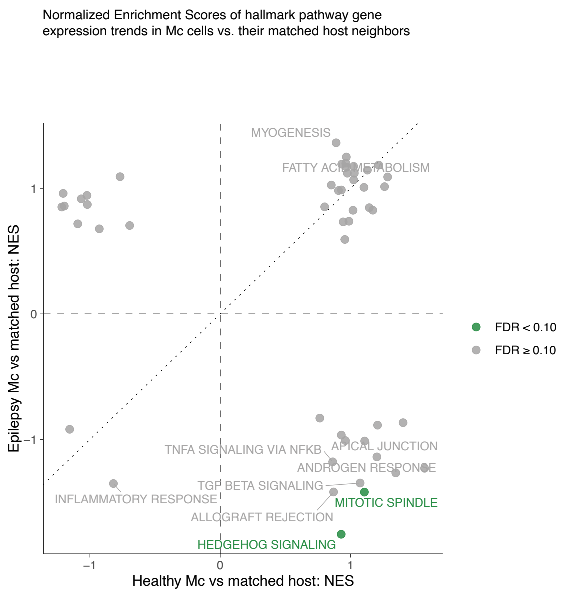

### SUPPLEMENTAL FIGURE 6

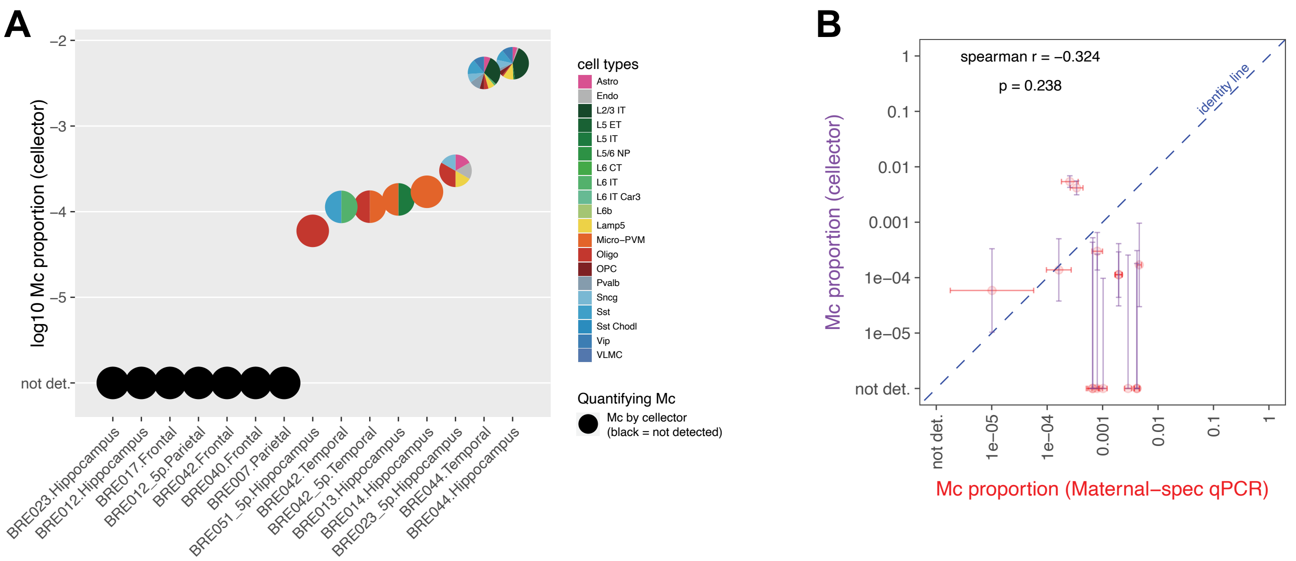

### SUPPLEMENTAL FIGURE 7

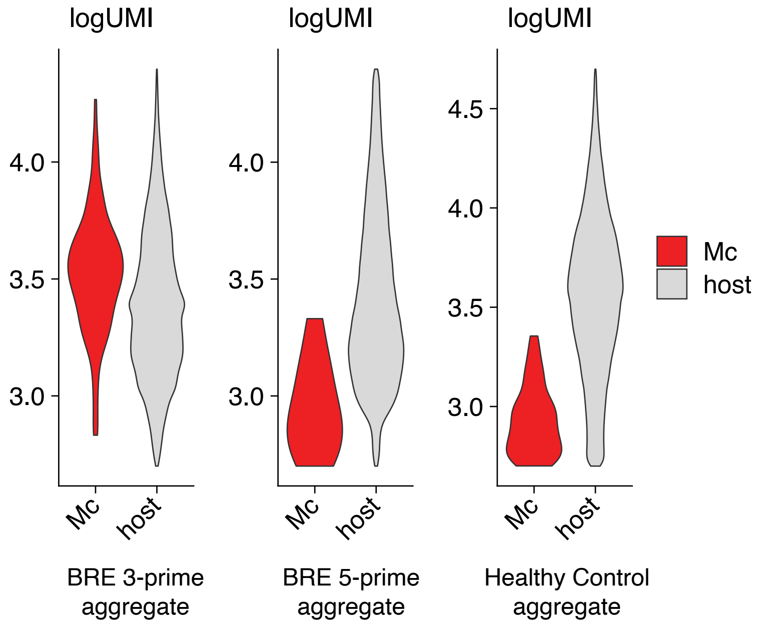
